# Proteins with proximal-distal asymmetries in axoneme localisation control flagellum beat frequency

**DOI:** 10.1101/2024.05.08.593170

**Authors:** Cecile Fort, Benjamin J. Walker, Lore Baert, Richard John Wheeler

**Affiliations:** Medawar Building for Pathogen Research, Nuffield Department of Medicine, University of Oxford, Oxford, UK; Department of Mathematical Sciences, University of Bath, Claverton Down, Bath, BA2 7AY, United Kingdom; Department of Mathematics, University College London, Gordon Street, London, WC1H 0AY, United Kingdom; Current acress: Swiss Tropical and Public Health Institute, University of Basel, Basel, Switzerland

**Keywords:** cilia, flagella, asymmetry, outer dynein arms, docking complex, flagellar beat

## Abstract

The 9+2 microtubule-based axoneme within motile flagella is well known for its symmetry. However, examples of asymmetric structures and proteins asymmetrically positioned within the 9+2 axoneme architecture have been identified in multiple different organisms, particularly involving the inner or outer dynein arms, with a range of functions. Here, mapped, genome-wide, conserved proximal-distal asymmetries in the uniflagellate trypanosomatid eukaryotic parasites. Building on the genome-wide localisation screen in *Trypanosoma brucei* we identified conserved proteins with an analogous asymmetric localisation in the related parasite *Leishmania mexicana*. Using deletion mutants, we map which are necessary for normal cell swimming, flagellum beat parameters and axoneme ultrastructure, and using combinatorial endogenous fluorescent tagging and deletion, map co-dependencies for assembly into their normal asymmetric localisation. This revealed 15 proteins, 8 known and 7 novels, with a conserved proximal or distal axoneme-specific localisation. Most were outer dynein arm associated, and showed that there are at least two distinct classes of proximal-distal asymmetry – one dependent on the docking complex, and one independent. Many were necessary for normal frequency of the tip-to-base symmetric flagellar waveform, and our comprehensive mapping reveals unexpected contribution of proximal-specific axoneme components to frequency of distal waveform initiation.

## Introduction

Flagella and motile cilia are microtubule-based organelles found across diverse eukaryotic lineages used for motility. They share a highly symmetric core architecture, known as the axoneme, based on nine doublet microtubules around a central pair of singlet microtubules. However, asymmetries in the distribution of axoneme-decorating proteins are emerging as being important for the function of cilia and flagella^1^.

Flagella undergo different waveforms specific to their function: typically, a planar symmetric near-sinusoid flagellar-type beat (e.g. human sperm) or a planar asymmetric ciliary-type beat (e.g. ciliated epithelia). Some flagella can switch waveform type, with *Chlamydomonas* adopting an asymmetric beat for normal swimming and a symmetric beat during a bright-light phototactic response^2–4^, in this case maintaining a base-to-tip direction of waveform propagation. Other examples can also switch waveform direction between base-to-tip and tip-to-base (e.g. *Leishmania*)^5,6^. *Leishmania* are one of the trypanosomatid parasites, a family which also includes *Trypanosoma brucei*. Due to tractable reverse genetics, these uniflagellate human pathogens are powerful model systems for analysing flagellar biology^7,8^. Trypanosomatid parasite flagellum-driven motility is necessary for normal life cycle progression^9,10^.

The flagellar beat is generated by the action of dynein complexes attached to the axoneme. The outer dynein arms (ODAs) attached to the nine doublets every 24 nm and are canonically viewed as being the primary driver of the flagellar beat^11^. Contrastingly, there are multiple different inner dynein arm (IDA) complexes attached in a larger 96 nm^12^ repeating unit, viewed as necessary for controlling the flagellar beat waveform rather than generating it^11^. However, recently, we showed that the preferred direction of waveform propagation in trypanosomatids is associated with a linear proximal-distal asymmetry in the ODAs^1^. We previously identified two paralogues of the ODA-docking complex (DC) heterodimer^13^, one heterodimer specific to the proximal and one to the distal axoneme^1^. Deletion of dDCs lead to loss of the distal ODAs, which caused a switch from the normal tip-to-base symmetric beat to the rarer base-to-tip asymmetric beat. While this precise DC-dependent asymmetry appears specific to the trypanosomatids, analogous DC or ODA asymmetries are found in diverse eukaryotes, including the unicellular parasite *Giardia*, the green alga *Chlamydomonas* and Humans^1^

This is not the only potentially conserved proximal-distal asymmetry. ARL13B is a well-characterised conserved marker of cilia and flagella necessary for normal ciliary / flagellar length^14–21^. However, in some tissues, including mouse oviduct and tracheal tissue, ARL13B is enriched in the proximal axoneme^22^. A similar proximal localisation is seen in *T. bruce*i^21^. Phosphodiesterases (PDEs) have also been identified as enriched in the distal flagellum: PDEA in *L. mexicana*^9^ and PDEB in *T. brucei* ^23,24^. This suggests complexity in proximal-distal axoneme composition beyond the ODAs.

Cyclic AMP (cAMP) and calcium ion (Ca^2+^) signalling are often implicated in the control of beat type to control cell motility^25–31^. For example, Ca^2+^ signalling is involved in the phototactic response of *Chlamydomonas*^32–34^. Trypanosomatid parasites are no exception: Ca^2+^ and cAMP alter beat type of demembranated reactivated *Leishmania* axonemes^35^ and there are likely links to asymmetrically positioned proteins. FLAM6 in *T. brucei*^36^, has a proximal DC-like localisation and has a predicted cAMP binding domain, flagellar cAMP signalling by PDEB is necessary for productive infection of the *T. brucei* insect vector^37^, and we previously identified LC4-like which has a predicted Ca^2+^ binding domain as a distal axoneme specific ODA-associated protein necessary for normal beat frequency in *T. brucei* and *Leishmania*^1^. Overall, this suggests interplay of signalling and asymmetric protein distribution.

It is becoming clear that complex proximal-distal asymmetries exist in the axoneme, and there are hints that proteins with a proximal-distal asymmetry in axoneme localisation may function in flagellum beat control. We therefore sought to map, genome-wide, all conserved proximal-distal asymmetries in a model flagellum. Using genome-wide subcellular protein localisations in *T. brucei,* we identified all proximal- and distal-specific proteins and identified those with a *L. mexicana* ortholog and an analogous sub-axonemal localisation. We show that *L. mexicana* have at least two proximal and distal asymmetries. A cohort of proximal-distal proteins are DC dependent or/and ODA heavy chain associated, including a new paralagous pair of DC-associated proteins where one is proximal and the other distal. Finally, we demonstrated that a subset of proximal and distal-specific proteins are necessary for normal control of flagellum beating, including, surprisingly, that proximal-specific proteins can influence frequency of waveforms starting at the distal tip of the flagellum.

## Results

### Fifteen proteins have proximal-distal asymmetry conserved between *T. brucei* and *L. mexicana*

To comprehensively identify proximal- and distal-specific proteins that may be responsible for flagellum beat control, we used the TrypTag (genome-wide subcellular protein localisation in *Trypanosoma brucei*) dataset^38^. Through a manual survey of proteins annotated with a flagellum or axoneme localisation, we identified 55 proximal-or distal-specific flagellar proteins. We reasoned that those with an *L. mexicana* ortholog that also has a proximal or distal-specific localisation are most likely to be functionally important. Of these 55, we therefore retained those with an *L. mexicana* ortholog (Table S1, S2, Figure S1, Figure 1) and did not further analyse those that lack an ortholog (Table S3, Figure S2, S3). To determine if proximal- or distal-specific localisation was conserved between *T. brucei* and *L. mexicana,* we tagged the *L. mexicana* proteins with mNeonGreen (mNG) at the C or N terminus at their endogenous loci (Figure 1). Overall, ∼50% (25/55) of *T. brucei* proximal or distal-specific proteins have a *L. mexicana* ortholog, of which ∼60% (15/25) have a comparable asymmetric localisation, leaving 10 proteins for detailed analysis.

**Figure 1.**
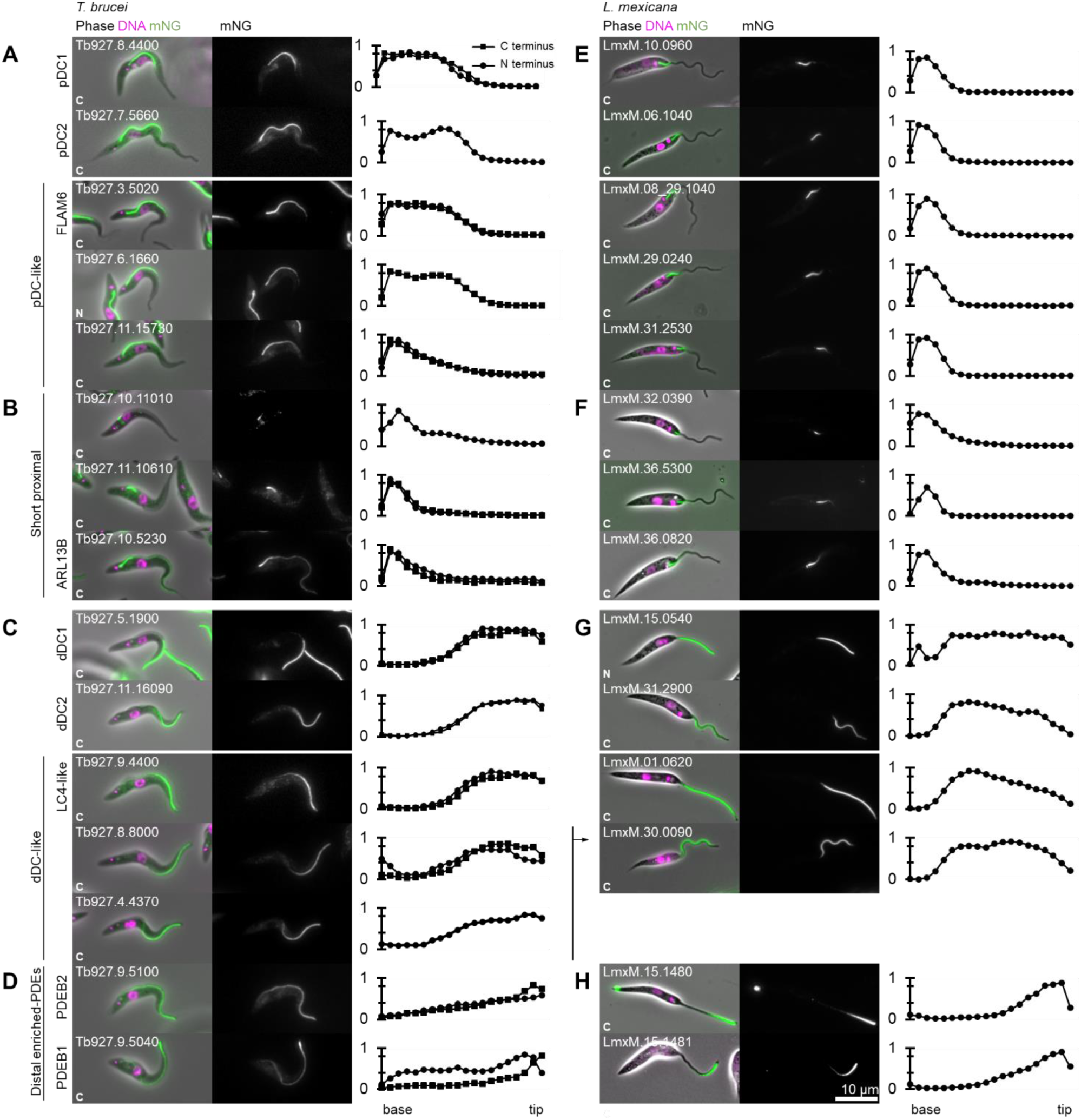
Comparative analysis of *T. brucei* proximal and distal-specific axoneme proteins in *L. mexicana* revealed 5 localisation groups. Quantitative analysis of fluorescence signal distribution along the axoneme from proximal and distal specific axoneme proteins endogenously tagged with mNG at the N and/or C terminus. Each row corresponds to a *T. brucei* protein and its *L. mexicana* ortholog. In the first and third columns, an example of *T. brucei* and *L. mexicana* cells, respectively. Phase contrast (grey), DNA (Hoechst 33342, magenta) and mNG (green) overlay (left) and mNG fluorescence (right) are shown. In the second and the fourth columns, graphs representing the mNG fluorescence signal intensity along the axoneme, from the base to tip. Data points represent the mean of *n* = 15 axonemes in 1K1N cells, normalised by maximum signal intensity per cell. From this analysis in *T. brucei* and *L. mexicana,* we infer 4 protein localisation groups: **A,E.** pDC-like. **B,F.** short proximal. **C,G.** dDC-like. **D-H.** Distally enriched proteins, only observed for PDEs.

Based on the sub-flagella localisation in these two species, we asked whether there are additional characteristic asymmetries in protein distribution distinct from the previously-characterised pDC/dDC asymmetry^1^. For each *T. brucei* and *L. mexicana* cell line, we measured the fluorescence signal distribution along the axoneme (Figure 1). pDC1 and pDC2 and dDC1 and dDC2 regions have a characteristic ∼50%-50% proximal-distal distribution in *T. brucei* and ∼20%-80% proximal-distal in *L. mexicana*^1^ (Figure 1A). We identified 3 proteins with a localisation similar to pDCs and 2 similar to dDCs in both species. We named these “pDC-like” and “dDC-like” localisations (Figure 1A, E and Figure 1C,G). This included a paralogous pair, one with a pDC-like and one with a dDC-like localisation. In *L. mexicana* LmxM.29.0240 (pDC-like) and LmxM.30.0090 (dDC-like). In *T. brucei* Tb927.6.1660 (pDC-like) and Tb927.8.8000/Tb927.4.4370 (dDC-like), where the latter is a more recent gene duplication due to the partial chromosome 4/8 duplication^39^.

A subset of proximal-specific proteins had a short ∼20% proximal signal in both *T. brucei* and *L. mexicana*. We named this the “short proximal” localisation (Figure 1B, F). This group included ARL13B which also had a weak fluorescent signal along entire the flagellum, as previously described in *T. brucei* procyclic forms^21^. Two proteins localised along the length of the flagellum with stronger signal towards the distal tip in *T. brucei.* These were PDEB1 and 2, replicating the previously described localisation in *T. brucei* bloodstream forms^24^, and we observed a corresponding enrichment in the distal ∼30% of the *L. mexicana* flagellum. We named this localisation group “distal enriched PDEs” (Figure 1D, H).This meta-analysis of two related species with differing morphologies suggests at least two types (DC-dependent and independent) of proximal and distal asymmetries.

### Deletion of novel asymmetrically distributed proteins has little impact on axonemal structure

Deletion of axoneme proteins can cause disruption of axonemal structure, therefore we generated deletions of each protein with proximal or distal-specific localisations in *L. mexicana.* Complete loss of the respective open reading frames (ORFs) was confirmed using diagnostic PCR from purified genomic DNA from each cell line (Figure S4). We were able to generate all deletion mutants by replacement of both alleles with drug-selectable markers, indicating that none of these proteins are vital. Flagellum length was near normal in all except one deletion mutant: ΔARL13B had shorter flagella (Figure 2A,B) as previously observed in *T. brucei*^14^ along with many other species^14,40^. This confirms ARL13B is functioning as expected in *Leishmania*, consistent with ARL13B mutations in humans causing Joubert syndrome ciliopathy ^20,41,42^ All but one of the proteins are therefore axoneme cytoskeleton components rather than factors necessary for axoneme assembly.

**Figure 2.**
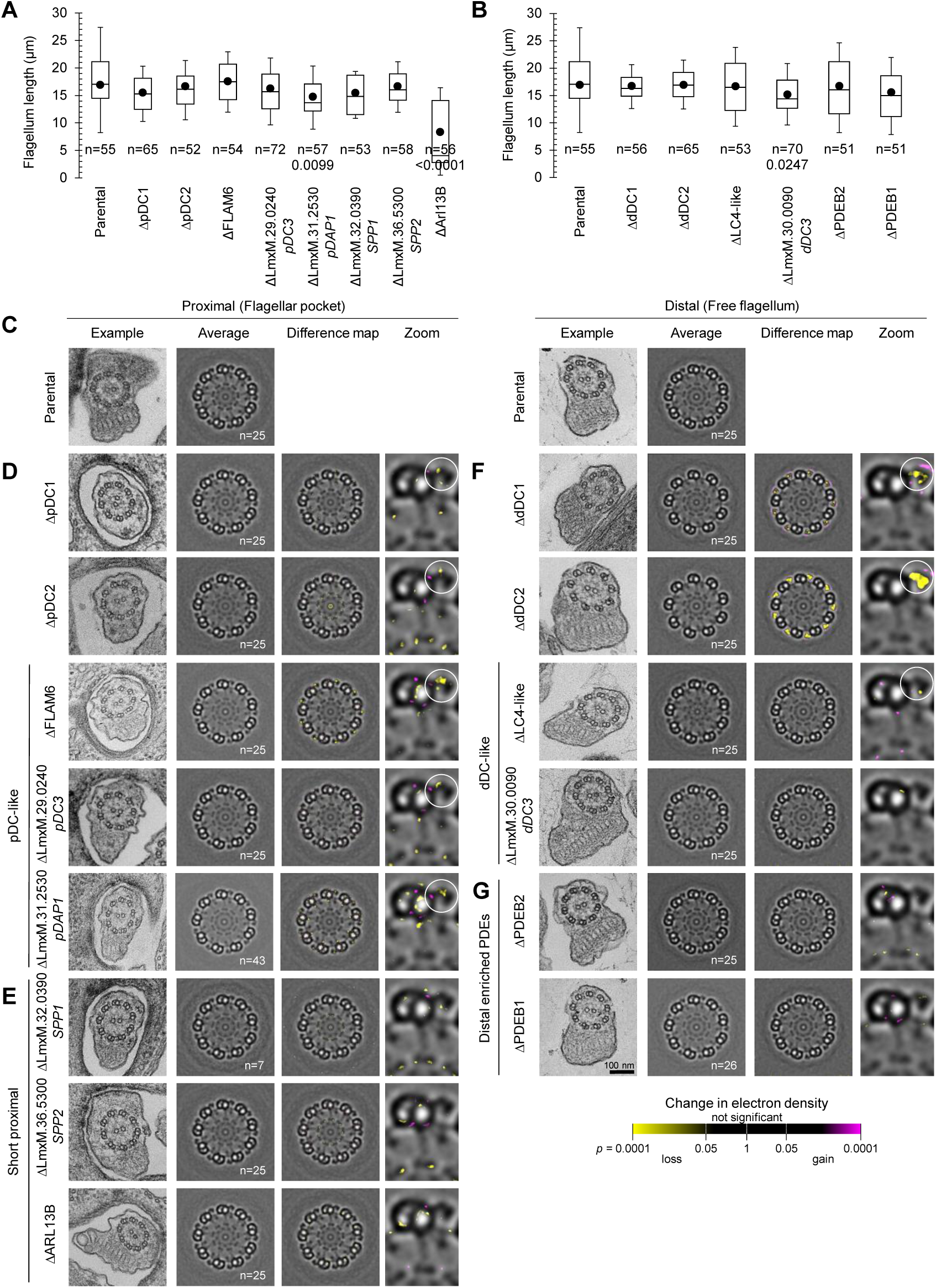
Deletion of proteins with pDC and dDC-like localisations cause minor changes in axonemal structure. **A-B.** Box and whisker plot of flagellum length in deletion mutants. Points represent the mean; box and whiskers represent the quartile ranges and the 5^th^ and 95^th^ percentile. n indicates the number of cells. Statistically significant differences (p<0.05, two-tailed T test) are indicated. **C-G.** Thin section electron microscopy of axoneme structure. The first column of each shows one representative axoneme cross-section in the flagellar pocket or in the free flagellum; the second column shows an averaged axoneme structure, in which axoneme cross-sections have had perspective deviation from circularity corrected (n indicates the number of axonemes used); the third column shows the result of ninefold rotational averaging and averaging across multiple axonemes; the fourth column shows a zoomed view of one microtubule doublet. Electron micrographs of transverse sections of axonemes in **C.** Parental cell line and **D.** Proximal pDC-like, **E.** Short proximal and **F.** distal dDC-like and **G.** Distal enriched PDEs proteins mutants. There is a specific loss in outer dynein arm (ODA) structure in dDC1/dDC2 images.

To identify any critical role in forming the axoneme ultrastructure, we analysed the flagellum structure of the deletion mutants by thin section transmission electron microscopy. In trypanosomatids, the flagellum protrudes from an invagination at the flagellar base – the flagellar pocket. The short pDC axoneme region of *L. mexicana* means that all cross-sections of flagella protruding through a flagellar pocket will be within the pDC-like and proximal-short axoneme regions (Figure 2C-E), while cross-sections through free flagella will largely (∼90%) be in the dDC-like axoneme region and ∼40% will be in the distal enriched PDEs region (Figure 2C, 2F,G). In both flagellar pocket and free flagellum cross-sections, no large defects in the axoneme organisation were visible: the ninefold symmetry of the outer doublets and the central pair was unchanged (Figure 2C-G, first column). As might be expected for proteins restricted to a small axoneme sub-domain, no proteins restricted to the pDC or dDC regions were necessary for normal overall axoneme organisation.

Next, we analysed changes in electron density in the flagellar-pocket and free-flagellum axoneme cross sections in these deletion mutants to identify subtle changes to axoneme structure. Making use of the ninefold radial symmetry of the outer dynein arms, we first perspective-corrected then ninefold rotationally averaged the axoneme cross-section images^43,44^. Subsequently, we used 25 rotationally averaged axoneme images to generate average flagellar pocket and free flagellum axoneme electron density maps and perform statistical comparisons to detect changes in electron density from the parental cell line (Figure 2). We confirmed the validity of this methodology by applying it to ΔpDC1, ΔpDC2, ΔdDC1 and ΔdDC2. Both dDCs are necessary for distal (but not proximal) ODA assembly. Contrastingly, neither pDC are necessary for proximal ODA assembly, as dDCs relocalise to fill the entire axoneme^1^. Concordantly, *Δ*pDC1 and *Δ*pDC2 had changes in electron density in the ODAs and at the point of attachment of the ODAs to the outer doublets in flagellar-pocket flagellar cross-sections (Figure 2D), while *Δ*dDC1 and *Δ*dDC2 had a large loss of electron density in the ODAs in free-flagellum flagellar cross-sections (Figure 2F).

We predicted that proteins with a pDC or dDC-like localisation will be associated with the DC/ODAs, and any electron density change will be in that axoneme region. In flagellar-pocket axoneme cross-sections, *Δ*LmxM.29.0240 had a small loss of electron density at the base of the ODAs (Figure 2D) and both *Δ*FLAM6 ^36,45^ and *Δ*LmxM.31.2530, are both predicted to have a cyclic nucleotide binding domain, had a clear loss of electron density within the ODAs.

Contrastingly, deletion mutants of proteins with a short proximal localisation (*Δ*LmxM.32.0390, *Δ*LmxM.36.5300, *Δ*ARL13B) had small or undetectable change in ODA electron density. These data were, however, relatively noisy, with changes in electron density around the central microtubule doublet and loss of electron density at the radial spoke head in several deletion mutants. As these included *Δ*pDC2, we believe these noisy observations are spurious, perhaps due to small variability in doublet position (Figure 2D, 2E). For proteins with a dDC-like localisation, in free-flagellum axoneme cross-sections, *Δ*LC4-like had a loss of electron density in the ODAs while *Δ*LmxM.30.0090 had no detectable change (Figure 2F). For the distal enriched PDEs, neither showed any detectable change (Figure 2G). Overall, the most significant changes in axoneme electron density were ODA-associated and for proteins with a pDC or dDC-like localisation. This suggests ODAs possess the primary structural asymmetries, although our method is only sensitive to proximal-distal, rather than doublet-to-doublet, asymmetries.

### There are two mechanisms for proximal and two for distal-specific localisations

To build direct evidence for whether proximal or distal-specific proteins are dependent on the DC asymmetry, we tested whether their localisation is dependent on DC proteins. First, for proteins with a pDC-like or short proximal localisation, we deleted pDC1 in the respective cell lines expressing the tagged protein and the successful deletion were confirmed by diagnostic PCRs (Figure S5). As previously observed^1^, this causes loss of pDC2::mNG proximal flagellum signal, as pDC1 and 2 are co-dependent for their normal localisation (Figure 3A). ΔpDC1 also caused loss of proximal axoneme signal for all proteins normally with a pDC-like localisation (FLAM6::mNG, LmxM.29.0240::mNG and LmxM.31.2530::mNG) (Figure 3A). None of the proteins normally with a short proximal localisation (LmxM.32.0390::mNG, LmxM.36.5300::mNG and ARL13B) lost proximal axoneme signal on pDC1 deletion (Figure 3B). To confirm this result, we deleted dDC2 in these tagged lines. Again, as previously observed, this causes extension of pDC1::mNG and pDC2::mNG signal to fill ∼30% of the flagellum (Figure 3A)^1^. All three proteins with a normally pDC-like localisation behaved similarly on dDC2 deletion (Figure 3A), while proteins normally with a short proximal localisation were unaffected (Figure 3B). Our classification of proteins by localisation into pDC-like and short proximal therefore represents a dependency on the DC asymmetry for their localisation.

**Figure 3.**
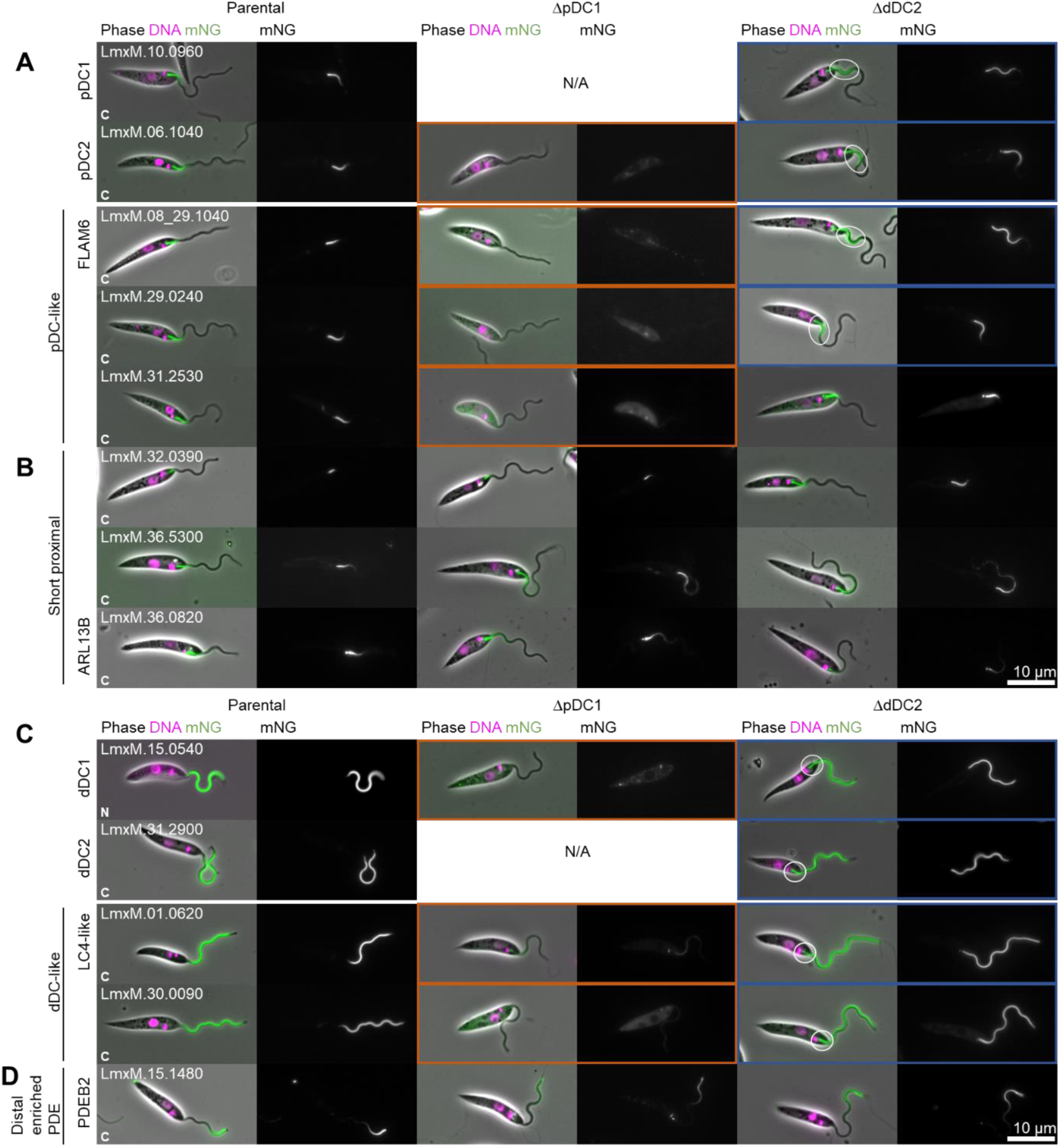
Proximal and distal-specific localisations can be either DC asymmetry dependent or independent. Protein localisation changes on pDC1 or dDC2 deletion. First column: micrographs of *L. mexicana* cell line expressing tagged proximal and distal proteins. Second column: after deletion of both alleles of pDC1. Third column: after deletion of dDC2. Phase contrast (grey), DNA (Hoeschst 33342, magenta) and mNG (green) overlay and mNG fluorescence are shown. Tagged proteins are grouped by localisation. **A.** pDC-like. **B.** Short proximal. **C.** dDC-like, **D.** Distal enriched PDEs. Cell lines where the protein fails to localise to the axoneme are outlined in orange, and cell lines where the localization is axonemal but changed are outlined in blue.

We next carried out the corresponding experiment for proteins with a distal-specific localisation. Here, dDC2 deletion causes loss of distal axoneme signal for dDC1::mNG and LC4-like::mNG (Figure 3C), as would be predicted from our previous work^1^, and LmxM.30.0090::mNG behaved similarly (Figure 3C). However, the localisation of PDEB2::mNG, a representative distal enriched PDEs, was unaffected (Figure 3D). pDC1 deletion causes expansion of signal dDC1::mNG, dDC2::mNG and LC4-like::mNG to fill the entire axoneme, again as expected^1^, and LmxM.30.0090::mNG behaves similarly (Figure 3C). PDEB2::mNG localisation was unaffected (Figure 3D). We have therefore accurately also identified the cohort of distal proteins dependent on the DC asymmetry. Overall, this conclusively shows that there are two mechanisms for proximal and distal-specific localisation occurring in trypanosomatid parasites.

### A subset of DC-dependent asymmetrically positioned proteins are ODA heavy chain associated

To dissect where proteins with pDC-like and dDC-like localisations sit within the ODA-DC complex proteins, we tested whether their localisation was dependent on ODA dynein heavy chains. However, first, we tested the co-dependency of ODA dynein heavy chains on their localisation. Similar to humans, trypanosomatids have two ODA dynein heavy chains, ODAα and ODAβ. These are orthologous to *C. reinhardtii* ODAγ and ODAβ respectively. In *C. renhardtii*, the ODA complex assembles in the cytoplasm prior to transport into the axoneme^46^, and mutation of either *C. reinhardtii* ODAγ or ODAβ prevents this process^47^ and *ODA4*^48^ have disrupted ODAγ or ODAβ respectively and completely lack ODAs). Thus, we expect *L. mexicana* ODAα and ODAβ to be co-dependent for normal localisation. Confirmation of ODAα and ODAβ deletion is shown in Figure S6A. As expected, deletion of ODAα in a cell line expressing ODAβ::mNG showed complete loss of ODAβ::mNG signal from the axoneme. However, surprisingly, deletion of ODAβ in a cell line expressing ODAα::mNG caused loss of ODAα::mNG signal from only the distal axoneme (Figure 4A).

**Figure 4.**
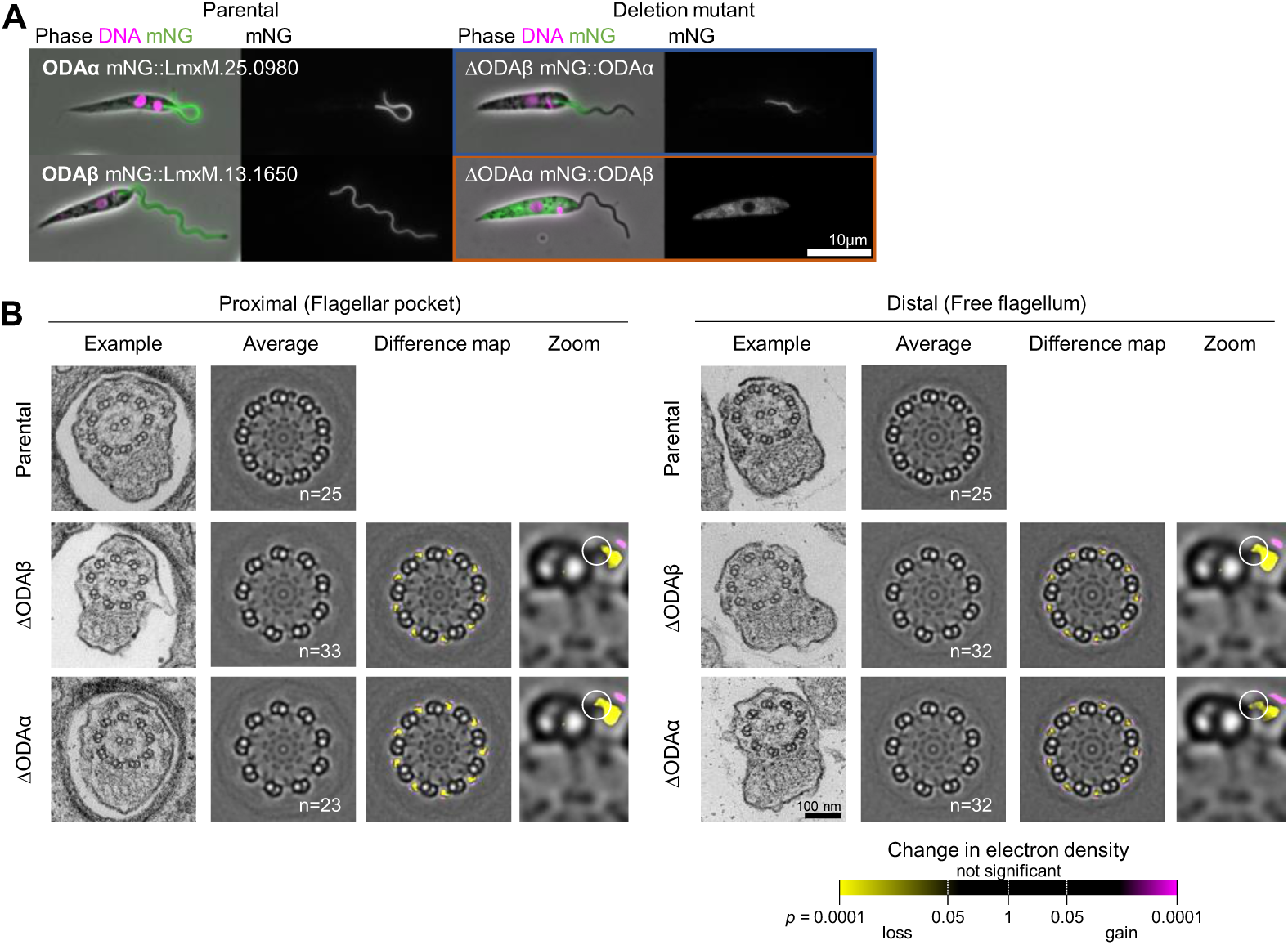
ODAβ deletion limits ODAα incorporation in the proximal axoneme while ODAβ requires ODAα for axonemal incorporation. **A.** First column: micrographs of *L. mexicana* cell line expressing ODAα or ODAβ tagged with mNG at the C terminus. Second column: before and after deletion of both alleles of ODAα or ODAβ. Phase contrast (grey), DNA (Hoeschst 33342, magenta) and mNG (green) overlay and mNG fluorescence are shown for. **B.** Ultrastructure changes upon ODA beta and ODA alpha deletion. First and second columns: one representative axoneme cross-section in the flagellar pocket and in the free flagella. Third and fourth columns: an averaged axoneme structure. (n indicates the number of axonemes used). Fifth and sixth columns: an electron density difference map, resulting from subtraction of deletion mutant average axoneme image from the parental cell line. Yellow indicates a loss of electron density in the deletion mutant. Cell lines where the protein fails to localise to the axoneme are outlined in orange, and cell lines where the localization is axonemal but changed are outlined in blue.

We confirmed this result by investigating the *Δ*ODAα and *Δ*ODAβ cell lines by thin section electron microscopy, investigating proximal (flagellar-pocket) and distal (free-flagellum) cross-sections in both deletion mutants (deletion validation is shown in Figure S6B). In *Δ*ODAα, ODAs were completely absent, while ODAs were only completely absent in the distal flagellum in *Δ*ODAβ. In the proximal flagellum (sections in the flagellar pocket), part of the ODA was present – it appeared that the innermost half of the ODA remained (Figure 4B). This is consistent with the expected position of ODAα as the innermost ODA dynein heavy chain, but not with ODA assembly requiring full cytoplasmic pre-assembly prior to flagellum import. Therefore, ODAs require ODAα but, unlike in *Chlamydomonas*, ODAs can incompletely assemble preferentially in the proximal axoneme in the absence of *Δ*ODAβ.

On the basis of this result, we analysed whether proximal or distal-specific proteins bind to the ODAs rather than the DC. For proximal proteins, this required deletion of ODAα (Figure 5A, validation of deletion Figure S7A), while for distal proteins we deleted ODAβ (Figure 5B, validation of deletion Figure S7B). As expected, pDC1::mNG, mNG::dDC1 and dDC2::mNG are not dependent on an ODA heavy chain for their normal localisation (Figure 4C, D). Among proximal proteins, FLAM6::mNG, LmxM.36.5300::mNG, ARL13B::mNG signal was lost on ODA heavy chain deletion, indicating that these are likely associated with the ODAs rather than the DC directly (Figure 5A). Among distal proteins, LC4-like::mNG and LmxM.30.0090::mNG localisation was altered on ODA heavy chain deletion, both with reduced signal and failure to localise all the way to the flagellar tip (Figure 5B).

**Figure 5.**
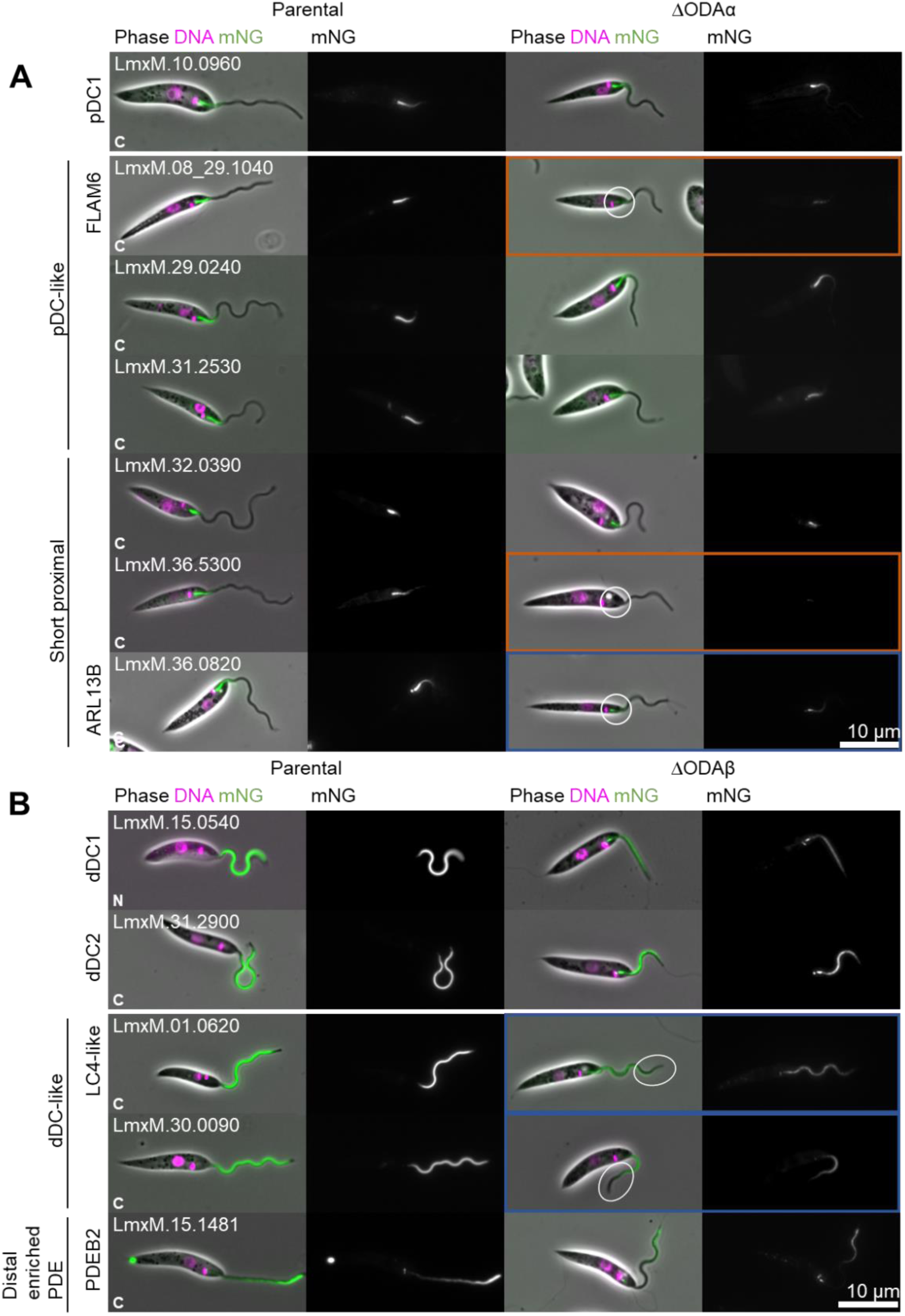
Some non-DC dependent localisations are ODA dynein heavy chain dependent and some DC dependent localisations are ODA heavy chain independent. **A.** Micrographs of *L. mexicana* cell line expressing proximal pDC-like and short proximal proteins and **B.** Distal dDC-like proteins tagged with mNG at the C terminus, before and after deletion of both alleles of ODAβ. Cell lines where the protein fails to localise to the axoneme are outlined in orange, and cell lines where the localization is axonemal but changed are outlined in blue.

Normal distal PDEB::mNG signal was ODA-independent. This reveals complexity: proteins can be proximal-specific and associated with the ODAs, with their proximal localisation either dependent (FLAM6, LmxM.36.5300) or independent (LmxM.36.0830) of the asymmetry of the DCs attaching the ODAs to the doublet. There are, therefore, two different mechanisms involved for proximal-distal asymmetry of ODA-associated proteins. The pDC-dependent and ODA heavy chain-independent localisation of LmxM.29.0240 and LmxM.31.2530 is also potentially indicative of a function assembling pDCs.

### LmxM.29.0240 is necessary for pDC assembly

To dissect which proteins are necessary for the DC assembly, we first deleted proteins with a pDC-like localisation in cells expressing pDC1::mNG localisation or deleting proteins with a dDC-like localisation in cells expressing dDC2::mNG (validation of cell lines in Figure S8). LC4-like and LmxM.30.0090 deletion did not affect the distal localisation of dDC2::mNG, although some accumulation of dDC2::mNG around the transition zone occurred following LmxM.30.0090 deletion (Figure 6B, validation of cell lines in Figure S8B). Therefore, these proteins are not necessary for dDC assembly. Contrastingly, LmxM.29.0240 was necessary for normal pDC1::mNG localisation, while FLAM6 and LmxM.31.2530 were not (Figure 6A, validation of cell lines in Figure S8A). LmxM.29.0240 is, therefore, acting as a pDC component necessary for pDC assembly; thus we named it pDC3.

**Figure 6.**
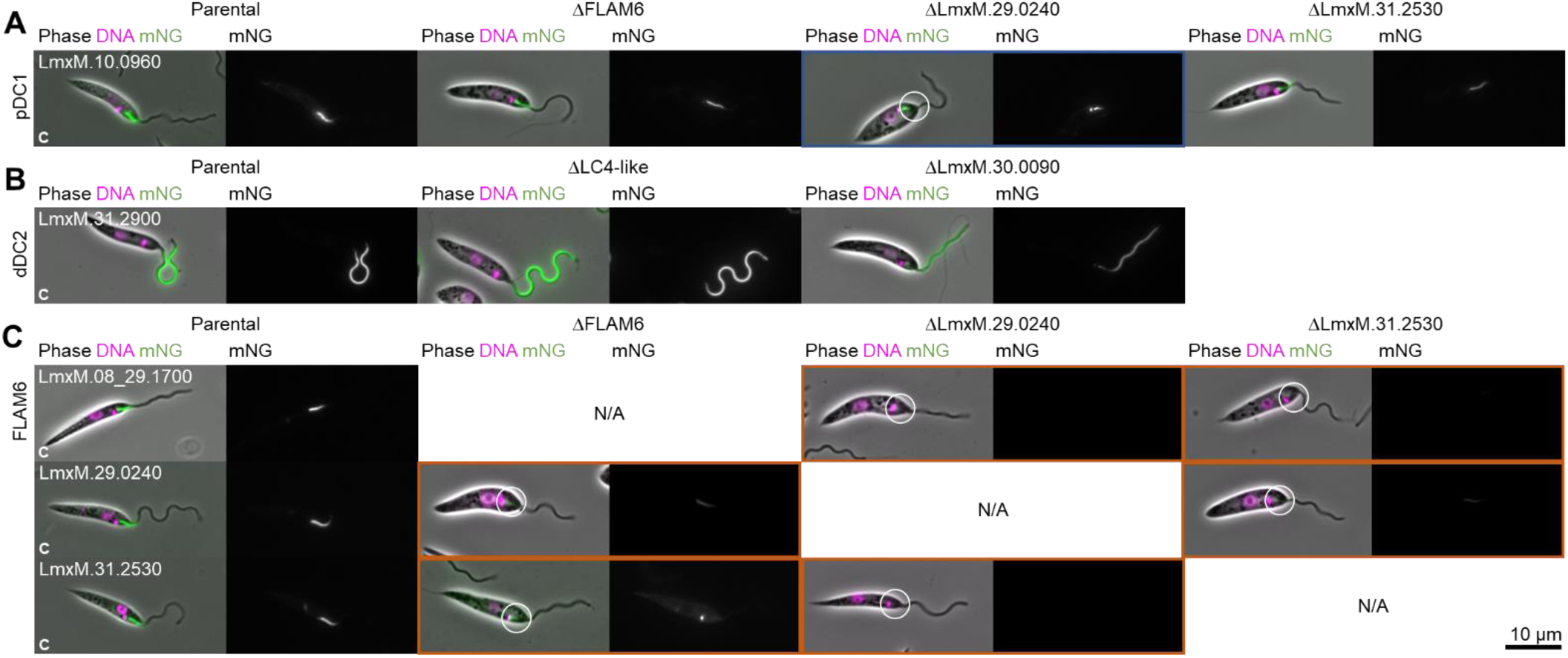
Localisation co-dependency among DCs and proteins with a DC-like localisation identifies an additional pDC component. **A.** Tagging of pDC1 and deletions of pDC-like proteins localisation. In the first column, micrographs of *L. mexicana* cell line expressing proteins tagged with mNG at the C terminus. In columns 2 to 4, micrographs after deletion of FLAM6, LmxM.29.0240 or LmxM.31.2530. **B.** Micrographs of *L. mexicana* cell line expressing dDC2 protein tagged with mNG at the C terminus, before and after deletion of both alleles of LC4-like and LmxM.30.0090. **C.** Combinatorial tagging and deletion of pDC1 or proteins with a pDC-like localisation. In the first column, micrographs of *L. mexicana* cell line expressing proteins tagged with mNG at the C terminus. In columns 2 to 4, micrographs after deletion of FLAM6, LmxM.29.0240 or LmxM.31.2530. Cell lines where the protein fails to localise to the axoneme are outlined in orange, and cell lines where the localization is axonemal but changed are outlined in blue.

Following our naming scheme for the proximal and distal DC1 and DC2 paralogous pairs, we named LmxM.30.0090 dDC3 – although it was not necessary for dDC assembly. To comprehensively test whether pDC3 is truly necessary for pDC assembly and map the co-dependencies of pDC-associated components, we used combinatorial tagging and deletion. pDC3 was necessary for the normal localisation of FLAM6::mNG and LmxM.31.2530::mNG in addition to pDC1::mNG (Figure 6C, validation of cell lines in Figure S8C). Normal LmxM.31.2530::mNG localisation was also dependent on FLAM6. pDC1, 2 and 3 therefore form the pDC, with FLAM6 and LmxM.31.2530 binding to the structure, LmxM.31.2530 in a FLAM6-dependent manner.

### Proximal and distal-specific proteins contribute to the control of the tip-to-base flagellum beat

Proteins that are not vital for flagellum assembly or normal ultrastructure but are well-conserved between related species are good candidates for flagellar beat regulators. To identify any correlation of beat regulation function with proximal or distal asymmetry, we carried out detailed cell swimming and flagellum beat analysis of all proximal and distal proteins not necessary for normal flagellum assembly. We carried out three analyses (Figure 7, S10): 1) Average cell swimming speed, in a deep volume and analysed away from the slide and coverslip to avoid surface interaction effects^16^; 2) Proportion of cells undergoing high frequency tip-to-base symmetric (flagellar-type), low frequency base-to-tip asymmetric (ciliary-type), low frequency aperiodic movement (static/uncoordinated) beats when in a thin volume between a slide and coverslip; 3) Beat waveform properties (amplitude, frequency and number of waves per flagellum) for cells in a thin volume between a slide and coverslip undergoing a tip-to-base symmetric waveform. For the latter, we noted that changes in mutants were dominated by change to beat frequency rather than amplitude, waves per flagellum, or the additional control measures (Figure S10).

**Figure 7.**
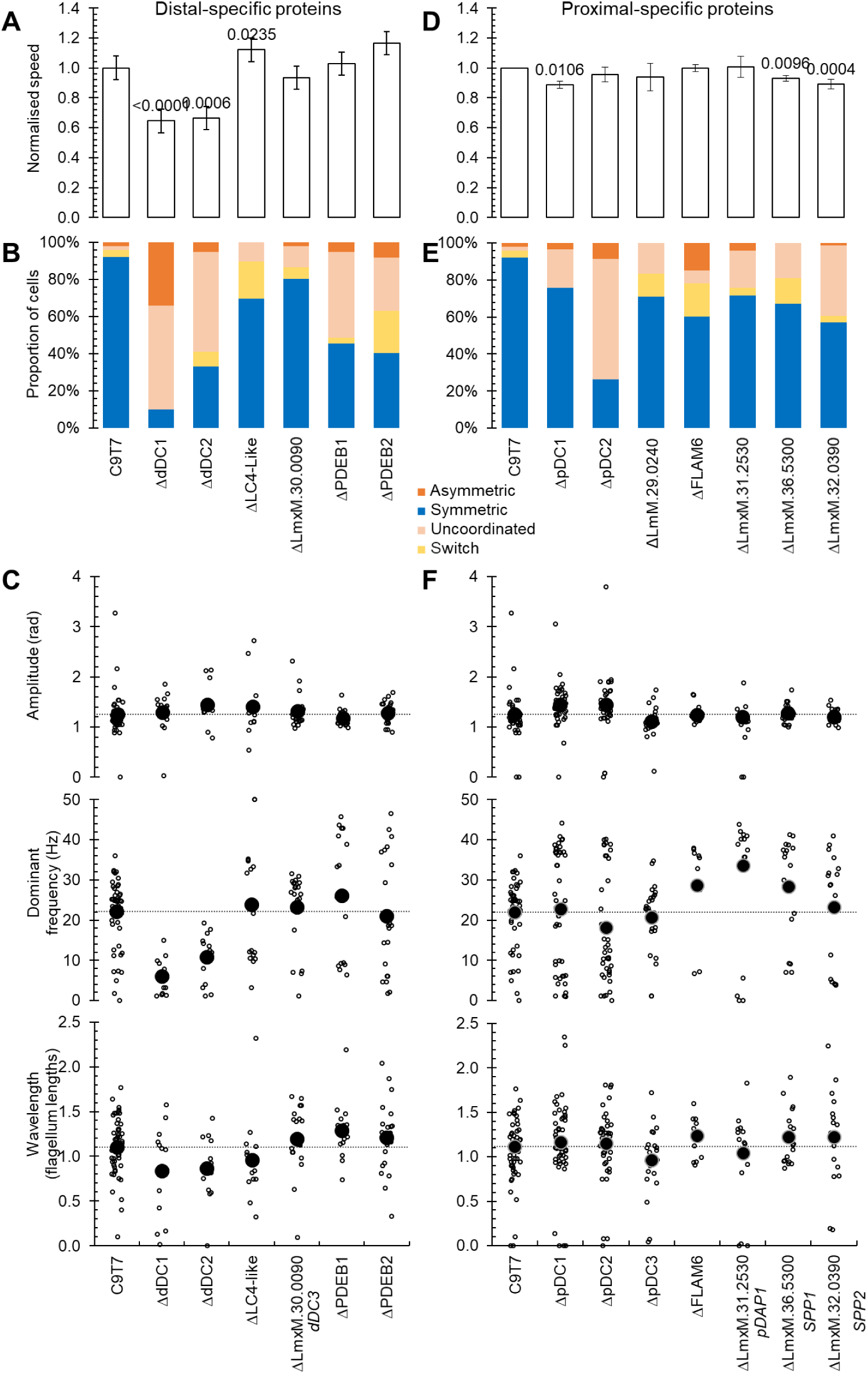
Distal and proximal proteins contribute to the control of flagellum beat and their frequency. **A,D.** Graphic representing the normalized swimming speed in parental cell line and all knockout mutants. Error bars represent the standard deviation of three replicates. p-values from Student’s t-test compared to the parental cell line. **B,E.** The proportion of cells undergoing different types of flagellar movement, comparing deletion mutants to the parental C9T7 cell line. **C,F.** Amplitude, dominant frequency and wavelength per flagellum in parental cell line and knockout distal mutants.

First, we consider proteins with a dDC-like localisation, where previous analyses are more comprehensive. As previously described, dDC1 and dDC2 deletion caused reduced swimming speed and reduced ability to carry out the normal tip-to-base symmetric beat (Figure 7A, B). When they did undergo a tip-to-base symmetric beat, it was much lower frequency. Again, as previously described, LC4-like deletion caused a small but significant increase in swimming speed, arising from a higher tip-to-base symmetric beat frequency. Deletion of the novel dDC-associated protein, LmxM.30.0090, had no significant effect on beat frequency, proportion of time spent undergoing a tip-to-base symmetric beat or beat frequency (Figure 7C). This confirmed the expected phenotypes but revealed little of novelty.

Deletion of either distal enriched PDEBs had little effect on swimming speed (Figure 7A) and reduced the number of cells able to undergo a tip-to-base symmetric beat (Figure 7B); however, those that did had a significantly increased beat frequency (Figure 7C). Distal enriched PDEs are therefore potential positive regulators of tip-to-base symmetric beats, but negative regulators of beat frequency.

Next, we considered proteins with a proximal localisation, seeing more surprising changes for both proteins with a pDC-like or short proximal localisation. In our previous work^1^, no large effect of pDC1 deletion on cell swimming speed was observed, thus was not analysed in detail. Here, consistent with this, pDC1, pDC2 and pDC3 deletion had little effect on swimming speed (Figure 7D). However, pDC2 and pDC3 deletion caused less tip-to-base symmetric beating, with a prominent increase in base-to-tip asymmetric beats upon pDC3 deletion (Figure 7E). pDC1 and pDC2 deletion changed tip-to-base symmetric beat frequency, giving a bimodal distribution of frequencies, i.e. cells tended to have either faster or slower beat frequency (Figure 7F). As pDC1, pDC2 and pDC3 are co-dependent for assembly of the pDC, we might expect these deletion mutants to have similar phenotypes. There was, however, mutant-to-mutant variability, but overall pDC proteins were necessary for a normal beat profile.

FLAM6, LmxM.31.2530 (pDC-like localisation), LmxM.31.0240 or LmxM.32.0390 (short proximal localisation) deletion all had little effect on swimming speed (Figure 7D) or proportion of time spent exhibiting different beat types (Figure 7E). However, all four had a large increase in beat frequency of tip-to-base symmetric beats (Figure 7F). Potentially, a proximal or distal-specific protein may have an effect on only the proximal or distal waveform. However, analysing frequency, amplitude and wavelength for the proximal and distal flagellum separately showed no significant difference between the two for any mutant with altered overall beat frequency (Figure S11). Therefore, surprisingly, proximal-specific axoneme proteins can be negative regulators of the flagellum-wide beat frequency of beats starting at the far end of the flagellum.

## Discussion

We have shown that proximal-distal asymmetry of axonemal organisation is a complex phenomenon. Analysis of genome-wide *T. brucei* data identified 55 examples of flagellar proximal-or distal-specific protein localisations. ∼50% (25/55) have orthologs among the trypanosomatid parasites and ∼30% (15/55) have an ortholog in *L. mexicana* with a comparable asymmetric localisation, summarised in Figure 8. Many of these are necessary for a normal flagellar beat, summarised in Table S4.

**Figure 8.**
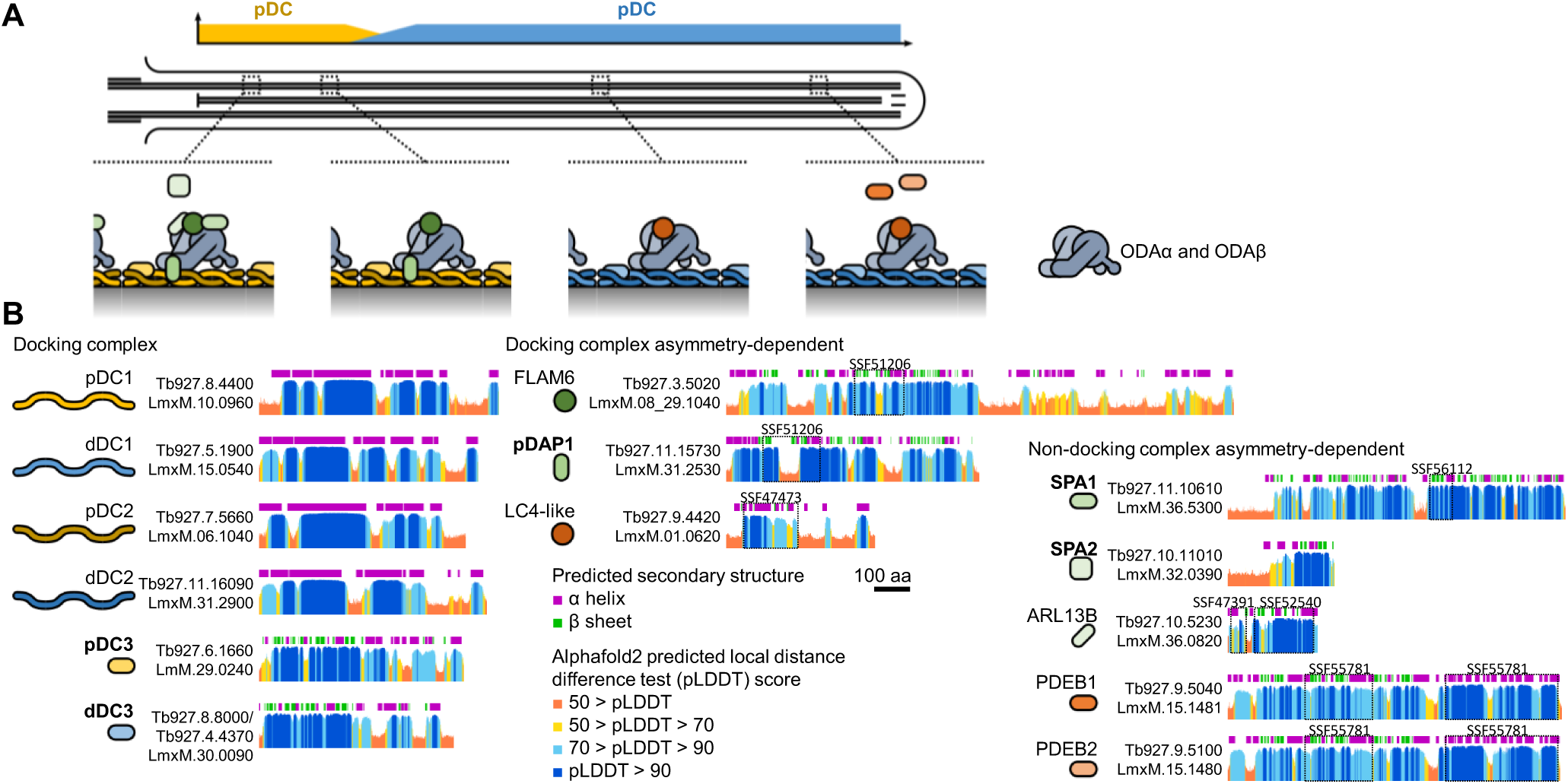
Summary of conserved proximal-distal asymmetry in the *Leishmania* and *Trypanosoma* flagellum. A. Cartoon summary of asymmetrically localised axonemal proteins, where proteins drawn overlapping indicate broadly summarises dependency for assembly. B. Summary of Alphafold2-predicted protein structure and predicted protein domains for the asymmetrically localised proteins in A.

Perhaps surprisingly, some aspects of proximal-distal asymmetry were evolutionarily well-conserved, while others appear to be recent innovations. This study identified several novel asymmetrically localised proteins whose localisation is conserved between *L. mexicana* and *T. brucei:* a paralogous pair associated with the proximal and distal docking complex, LmxM.29.0240 and LmxM.30.0090 which we name pDC3 and dDC3 respectively, a **p**roximal **D**C-**a**ssociated **p**rotein, LmxM.31.2530 which we name pDAP1, and two proteins which localise to a **s**hort **p**roximal **a**xoneme region, LmxM.36.5300 and LmxM.32.0390 which we name SPA1 and SPA2 respectively. We previously noted that ODA proximal-distal asymmetries occur across diverse eukaryotes^49–54^ although are not necessarily orthologous (i.e. not all involving a pDC and dDC pair of paralogous heterodimers) to *T. brucei* and *L. mexicana*^1^, and the majority of proteins we analysed in detail were ODA-associated (Figure 5), hinting at a tendency for proximal-distal asymmetries to generally arise in ODAs. Importantly, uncharacterised orthologs of pDC3/dDC3 and SPA2 are found in the genomes of several flagellate/motile ciliate species, including *Chlamydomonas reinhardii,* and Alphafold2 structure predictions of pDC3/dDC3, SPA1 and SPA2 predicted globular domains not identified by sequence-based protein domain detection (PFAM, Superfamily, etc.)^55–57^.

The proximal and distal flagellum appear, overall, similarly complex in terms of asymmetrically positioned molecular machinery. This applies to both genome-wide analysis of *T. brucei* and the set of proteins with an ortholog in *L. mexicana*. However, there was a large degree of apparently recent adaptation. Many asymmetrically positioned *T. brucei* proteins lacked an *L. mexicana* ortholog, and a further set had a whole flagellum rather than asymmetric localisation (Figure S1) – indicating either loss of asymmetry in *L. mexicana* or gain of asymmetry in *T. brucei*. Either way, this indicates notable adaptation within the different highly motile trypanosomatid parasite lineages on the time scale of hundreds of millions of years. It is an intriguing possibility that evolvability of flagellum beat control involves evolvability of asymmetric position within the flagellum.

We have confidently identified at least two distinct types of proximal and two types of distal asymmetry, indicating that there must be multiple mechanisms underlying their formation. Previously, we saw that the pDC/dDC asymmetry is likely achieved by competition for axoneme binding on top of IFT transport to form a proximal-distal concentration gradient, leading to mutually exclusive axoneme regions^1^. Interestingly, our data showed that OADα and OADβ were not fully co-dependent for assembly into the axoneme, unlike in *C. reinhardtii*, perhaps related to differences in DC biology compared to *C. reinhardtii.* Here, we comprehensively mapped the conserved proteins whose asymmetric localisation is dependent on the dDC/pDC asymmetry, finding pDC3, pDAP1 and FLAM6 as pDC-dependent and dDC3 and LC4-like as dDC-dependent. Interestingly, pDC3 is necessary for the correct assembly of the pDC1/2 heterodimer into the proximal axoneme while its paralog, dDC3, was not necessary for normal dDC1/2 distal axoneme localisation. Perhaps this is a key distinguishing feature of the proximal and distal docking complex.

The other axoneme asymmetries we identified do not seem to have the same mutual exclusivity as the pDC/dDC asymmetry: we identified novel short-proximal (SPA1 and SPA2) and short-distal regions (PDEB1 and PDEB2), but no corresponding conserved long-proximal and long-distal regions, respectively. A simple explanation for short-distal and short-proximal regions is limiting protein abundance, with their asymmetric localisation maintained by anterograde or retrograde transport by intraflagellar transport (IFT), respectively. Short-proximal localisation could also be explained by no active transport by IFT, only diffusion up the flagellum, and high binding affinity. The latter was previously proposed for ARL13B^21^ and we confirmed the proximal ARL13B localisation was indeed not linked with DC asymmetry, but its axonemal localisation was ODA-dependent. Contrastingly, distal PDEB localisation was not DC-dependent and not ODA-dependent.

There are, however, other potential mechanisms for generating asymmetry. One is tubulin post-translational modification, which could define specialised microtubule regions to which proteins could bind. Detyrosination is a well-characterised example in trypanosomatids^58–61^ and many other modifications could be infolved^62^. Alternatively, some axonemal regions may be defined by other intraflagellar or extraflagellar structures, like the intraflagellar para-axonemal structure called the paraflagellar rod (PFR)^63^ or the lateral attachment of the flagellum to the flagellar pocket neck by the flagellum attachment zone (FAZ)^64^. The PFR-free and FAZ structures broadly line up with the short-proximal region, but dependence of PFR or FAZ positioning on axoneme asymmetries is unlikely as we saw no obvious change to the PFR or FAZ by electron microscopy in the SPA1 and SPA2 deletion mutants.

Having mapped the conserved asymmetries in *L. mexicana* flagella, the critical question is the function of these asymmetrically positioned proteins. We showed that many asymmetrically positioned proteins are necessary for normal beating. However, we did not perturb asymmetry, thus do not have evidence that the asymmetric positioning is also necessary for normal beating. In general, deletion mutants had complex changes to the preferred mode of flagellum movement but, by focusing on only the tip-to-base waveform, we carried out systematic and comparable analysis of all deletion cell lines. This showed that deletion of dDC1/2 is an outlier phenotype. Total removal of distal ODAs by dDC1/2 deletion were the only mutants with a strong preference for a base-to-tip beat and, furthermore, on the rare occasion when these cells manage a tip-to-base beat, it was at low frequency. Other deletion mutants of asymmetrically positioned proteins either had little change to beat frequency or gave a significant population with increased tip-to-base beat frequency. Therefore, distal ODAs are needed for efficient distal waveform initiation, and asymmetrically positioned flagellar proteins are often negative regulators of beat frequency.

Interestingly PDEB deletion was previously shown to have no effect on *T. brucei* swimming speed^24^ and we saw no statistically significant effect of PDEB deletion on swimming speed in *Leishmania*, although PDEB2 deletion gave a small speed increase and PDEA deletion was previously shown to increase *Leishmania* swimming speed. While the effect on swimming speed may be marginal, variable^9^, or contingent on deleting both PDEBs, beat frequency was clearly and strongly affected by deletion of either PDEB. The *T. brucei* PDEB deletion mutant has defects in chemotaxis-like behaviour termed “social motility” and inability to progress through the tse-tse fly vector^23,24^. This likely is a result of changes to beat frequency control, and emphasises the importance of cAMP signalling to regulate flagella. We suspect the tendency for bimodal beat frequency, some increased and some decreased, is the result of an increase in beat frequency to the point that it cannot be stably maintained.

Roles in regulation of tip-to-base waveform frequency control are intuitive for distal-specific proteins, like PDEB. But deletion of the either FLAM6 or pDAP1, both pDC-associated, caused a significant increase in the tip-to-base beat frequency. How can removal of protein present only in a short proximal region of the flagellum increase beat frequency by promoting initiation of waveforms at the distal end of the flagellum?. We can consider a two key hypotheses: Their deletion may lead to a constant signal that increases the rate of tip-to-base waveform initiation thus increasesing frequency, or their deletion may allow faster transmission of individual signal that initiates individual tip-to-base waveforms in more rapid succession. The former seems unlikely – this requires proximal proteins to act as sinks for a molecule that acts as a constant negative signal. FLAM6 and pDAP1 are predicted to have cAMP binding domains, and cAMP can be a positive regulator of beating^26–28^. However, the predicted binding rather than enzymatic degradation represents a limited sequestration rather than a sink, making their deletion unlikely to have a large effect on flagellar cAMP concentration. The latter is possible, but only through limited mechanisms. Signal from the flagellar base would have to reach the tip in ∼25 ms to initiate successive tip-to-base waveforms at 40 Hz (a speed of ∼1,000 μm/s).

Diffusing or actively transported (*cf.* IFT train speed in trypanosomatids^65^) signalling molecules are far too slow; however, a mechanical signal like shear force on the outer doublet microtubules would be transmitted almost instantaneously. We suggest that these proximal-specific proteins act to modulate shear force generation as waveforms approach the base of the flagellum.

Overall, we have shown that there are multiple mechanisms for generating proximal-distal asymmetry in axoneme organisation, often involving the ODAs. Surprisingly, proximal-specific proteins can be necessary for normal beat frequency for flagellar waveforms starting at the distal end of the flagellum.

## Methods

### Parasite cell lines, maintenance and culturing

Cas9T7 *L. mexicana* derived from WHO strain MNYC/BZ/62/M379, expressing Cas9 and T7 RNA polymerase^8^ were grown in M199 (Life Technologies) supplemented with 2.2 g/L NaHCO_3_, 0.005% hemin, 40 mM HEPES HCl (pH 7.4) and 10% FCS. *L. mexicana* cultures were grown at 28°C. Culture density was maintained between 1 × 10^5^ and 1 × 10^7^ cells/mL for continued exponential population growth. Culture density was measured using a haemocytometer. Identity was confirmed by recent mRNA and genomic sequencing, and lack of mycoplasma contamination was confirmed by fluorescent microscopy with a Hoechst 33342 DNA stain.

### Protein sequence analysis

Tagging was carried out in *L. mexicana* and we considered genes for tagging if they had a syntenic ortholog in *T. brucei*. Ortholog proteins were identified by reciprocal best protein sequence search hits carried out using The BLAST Sequence Analysis Tool^66^.Genes were selected for tagging at N or C terminal tagging using TrypTag PCF protein localisation data available up to 12^th^ March 2018^67^ and TriTrypDB version 36. AlphaFold protein structure from genome in *T. brucei* and *L. mexicana* sequencing are done by the methods published by RJ Wheeler ^57^.

Domain identification, ortholog identification, structure prediction, etc. Cite tritrypdb, cite my alphafold.

### Genetic modifications

Constructs and sgRNA templates for endogenous mNG-tagging templates were generated by PCR as previously described^8^ and were transfected as previously described^68^. The pLrPOT series of vectors was used as PCR templates for generating tagging constructs^1^, specifically pLrPOT mNG Neo. Constructs and sgRNA templates for ORF deletion were generated by PCR and transfected as previously described, using pT Blast, pT Puro and pT Neo as templates^8^. Primers were designed using LeishGEdit (www.leishgedit.net/)^8^. Transfectants were selected with the necessary combination of 20 μg/mL puromycin dihydrochloride, 5 μg/mL blasticidin S hydrochloride and 40 μg/mL G418 disulfate.

### Diagnostic PCR for gene knockout validation

To verify loss of the target ORF in drug-resistant transfectants, a diagnostic PCR was performed by amplifying a short PCR product (100–300 bp) within the ORF of the target gene. We used a positive control of with genomic DNA from the parental cell line to confirm successful detection of the target ORF. Primers to amplify a short fragment of the PF16 (LmxM.20.1400) ORF was amplified as a technical control to confirm presence of genomic DNA from the knockout cell line. The PCR mixture for each reaction was: ≤100 ng of gDNA (as required) and 10 μM (1.25μL) each of the forward and reverse primers mixed with the FastGene Optima HotStart Ready Mix with dye (12.5 μL) (Nippon Genetics, [P8-0082]) up to 25 μL of PCR-grade water. The thermocycle: Step Initial denaturation at 95°C for 3 mins, then 30 cycles of 95°C for 15 seconds, 58°C for 15 seconds, 72°C for 30 seconds, then final elongation at 72°C for 1 min.

### Microscopy

*L. mexicana* expressing fluorescent fusion proteins were imaged live. Cells were washed three times by centrifugation at 800 g followed by resuspension in PBS. DNA was stained by including 10 μg/mL Hoechst 33342 in the second washing. Washed cells were settled on glass slides and were observed immediately. Widefield epifluorescence and phase-contrast images were captured using a Zeiss Axioimager.Z2 microscope with a 63×/1.40 numerical aperture (NA) oil immersion objective and a Hamamatsu ORCA-Flash4.0 camera. Cell morphology measurements were made in ImageJ ^69^.

### Motility Assays

For motility analysis in *L. mexicana,* swimming behaviours are analysed for cells in the exponential growth phase in normal culture medium essentially as previously described (Wheeler, 2017). For cell swimming analysis, a 25.6 s video at five frames/s under darkfield illumination was captured from 5 μL of cell culture in a 250 μm deep chamber using a Zeiss Axioimager.Z2 microscope with a 10×/0.3 NA objective and a Hamamatsu ORCA-Flash4.0 camera. Particle tracks were traced automatically, and mean cell speed, mean cell velocity and cell directionality (the ratio of velocity to speed) were calculated as previously described ^9^.

### Flagellum beat type analysis

To determine the proportion of cells in a population undergoing different beat types, 1 mL of exponential growth cells (between 1 × 10^6^ and 1 × 10^7^ cells/mL) was centrifuged for 5 min at 800 g. Between 700 and 950 μL (depending cell density) of supernatant was removed and cells were resuspended in M199 (300 to 500 μL). 1uL of 5 μm polysyrene beads diluted 1:100 in M199 (Sigma 79633) was added, which ensure a 5 μm sample depth. 1 μL of cell sample was added to the center of a 2 by 5 cm area marked with a hydrophobic pen on a slide, and a glass coverslip (1.0 thickness) added. Videomicrographs of swimming cells under phase contrast illumination were captured with an Andor Neo 5.5 camera at 200 frames/s for 0.5 sec, using a x20 NA 0.3 objective lens on a Zeiss Axioimager.Z2 inverted microscope. Cells with one flagellum (non-dividing) were manually classified into symmetrical tip-to-base (continuous or interrupted), asymmetric base-to-tip, wave type switch and static and uncoordinated.

### Flagellar beating analysis

Parental cell line and deletion cell lines were analysed by high-speed video microscopy. A 5 s video at 200 frames/s under phase-contrast illumination was captured from a thin film of cell culture between a slide and coverslip using ZeissAxioimager.Z2 microscope with a 100x/1.4 NA objective and an Andor Neo 5.5 camera. Flagellar beat behaviours for each cell lines were classified manually and only symmetrical tip-to-base waveforms were analysed for this study. Automated image analysis and flagellum tracking^70^ in ImageJ [version 1.52a] was used to digitise the flagellar waveforms of a target 25 cells (all at least 19), with a target of 950 (all at least 30 frames) per cell, manually excluding cells which swam out of focus or out of the frame during video capture. Digitised waveforms were screened based on the variation in measured flagellum length over each video (a proxy for consistency of digitisation), then smoothed in space and time with smoothing splines in MATLAB. Finally, waveforms were excluded if they were poorly approximated by a sinusoidal beat (wild-type *L. mexicana* has a sinusoidal tip-to-base beat), as measured by a least-squares fit. A range of beating characteristics were computed for each cell and all deletion cell lines were compared to the parental cell line. Full code for the analysis pipeline (including thresholds for exclusion) is available on request.

### Transmission electron microscopy

For transmission electron microscopy, *L. mexicana* were fixed directly in medium for 10 min at room temperature in 2.5% glutaraldehyde (glutaraldehyde 25% stock solution, EM grade, Electron Microscopy Sciences). Centrifugation was carried out at room temperature for 5 min at 16,000 g. The supernatant was discarded and the pellet was fixed in 2.5% glutaraldehyde and 4% PFA (16% stock solution, EM grade, Electron Microscopy Sciences in 0.1 M PIPES (pH 7.2)) for minimum 2h. Cells were embedded in 3% agarose and contrasted with OsO_4_ (1%) (osmium tetroxide 4% aqueous solution, Taab Laboratories Equipment) during 2 hours at 4°C. Cells are stained with 2% of Uranyl Acetate for 2h 4°C. After serial dehydration with ethanol solutions, samples were embedded in low-viscosity resin Agar 100 (Agar Scientific, UK) and left to polymerise at 60°C for 24 h. Ultrathin sections (90 nm thick) were collected on nickel grids using a Leica EM UC7 ultra microtome and stained with uranyl acetate (1%, w/v) (uranyl acetate dihydrate, Electron Microscopy Sciences) and Reynolds lead citrate^71^ (Lead nitrate (Thermofisher, L/1450), sodium citrate (Sigma, 71405) and sodium hydroxide (SLS, CHE3422). Observations were made on a Thermo Fisher Scientific Tecnai12 or JEOL 2100 Plus 200kV transmission electron microscope with a Gatan OneView camera.

### Ninefold rotational averaging of *L. mexicana* axonemes

For generation of averaged axoneme views, axoneme images were first perspective corrected to ensure circularity, followed by nine-fold rotational averaging as previously described^72^. 25 rotationally averaged axonemes were then aligned and averaged. Difference maps were generated by comparison with the average 25 rotationally averaged parental cell line, and per-pixel statistical significance of electron density changes calculated by Mann Whitney U test (with multiple-comparison correction for the number of pixels within the axoneme cross-section).

## Acknowledgments

We thank Dr Errin Johnson and Dr Charlotte Melia for technical assistance for transmission electron microscopy (Electron microscopy facility at Sir William Dunn School of Pathology, Oxford University, United Kingdom). This work was supported by a Wellcome Trust Sir Henry Dale Fellowship [211075/Z/18/Z] awarded to RJW. BJW is supported by the Royal Commission for the Exhibition of 1851.

## Supplemental Figures and Tables

**Figure S1.**
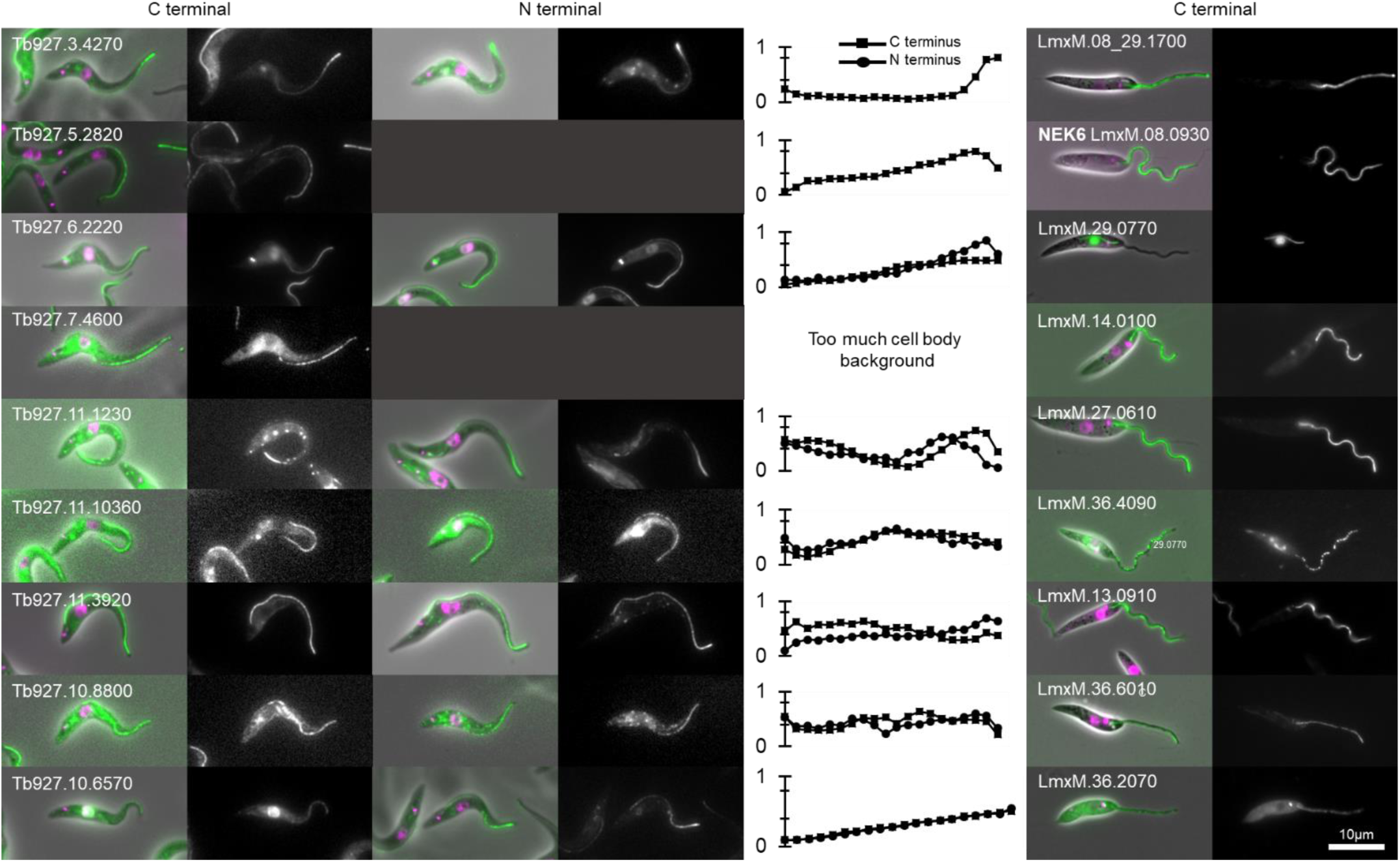
Proximal or distal axoneme-specific proteins in *T. brucei* where the *L. mexicana* ortholog did not have a proximal or distal-specific localisation. Localisation of proteins with a proximal or distal specific localisation in *T. brucei* but in *L. mexicana*. The first column shows widefield epifluorescence micrographs of endogenous mNG tagging at the N and C termini in *T. brucei*. Phase contrast (grey), DNA (Hoechst 33342, magenta) and mNG (green) overlay and mNG fluorescence are shown. The second column is a graph representation of the mNG fluorescent signal intensity along the axoneme from the base to the tip. Data points represent the mean of *n* = 15 axonemes in 1K1N cells, normalised by maximum signal intensity per cell. The third column shows widefield epifluorescence of endogenous tagging at the C terminus of the *L. mexicana* ortholog. These were categorised as not specific to the proximal or distal axoneme.

**Figure S2.**
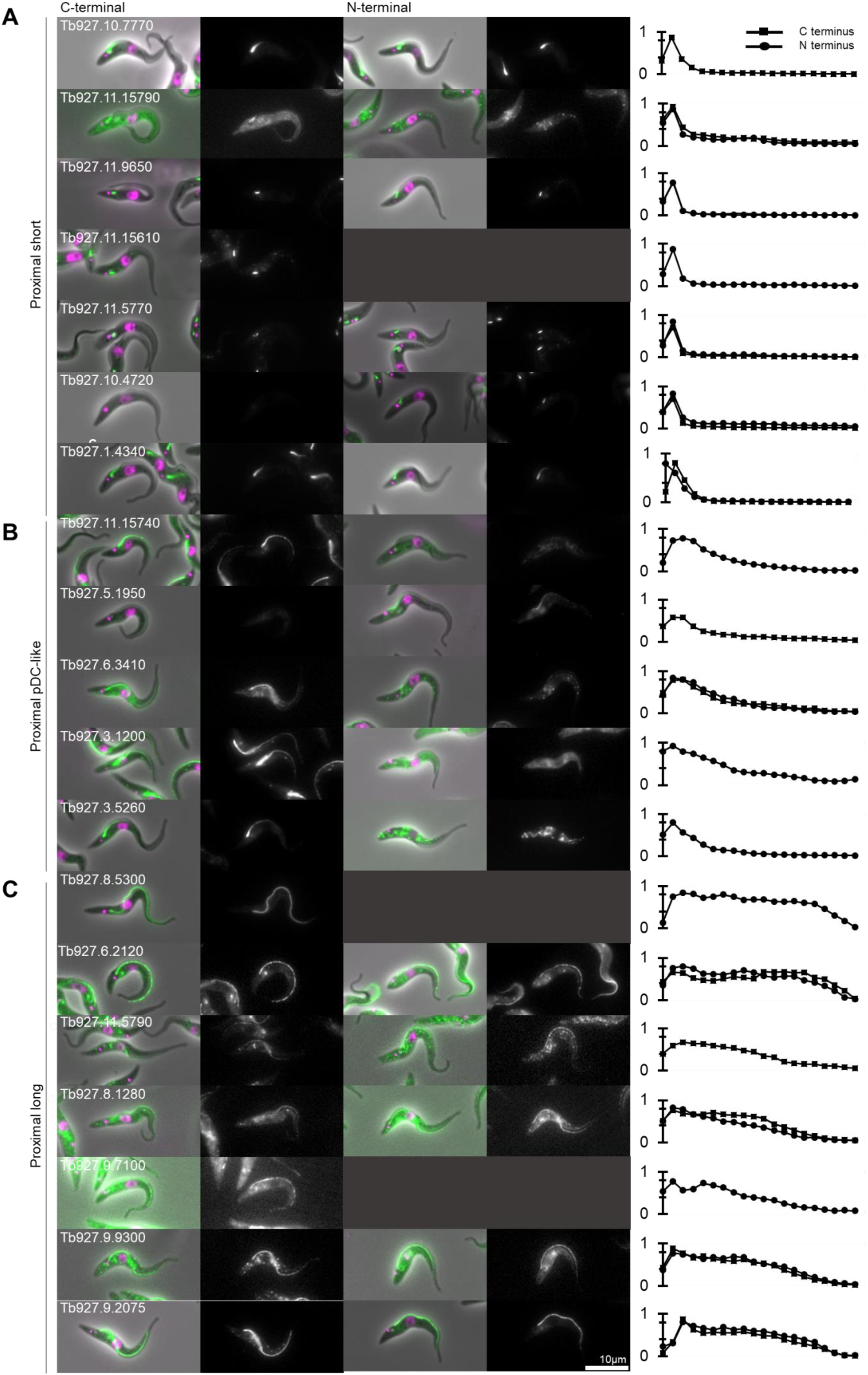
Proximal axoneme-specific proteins in *T. brucei* which lack a detectable *L. mexicana* ortholog. Quantitative analysis of fluorescence signal distribution along the axoneme from proximal specific axoneme proteins endogenously tagged with mNG at the N and/or C terminus. In the first column epifluorescence microscopy images of mNG fluorescence in example *T. brucei* cell. Phase contrast (grey), DNA (Hoechst 33342, magenta) and mNG (green) overlay (left) and mNG fluorescence (right) are shown. In the second column, graphs representing the mNG fluorescence signal intensity along the axoneme, from the base to tip. Data points represent the mean of *n* = 15 axonemes in 1K1N cells, normalised to maximum signal intensity per cell. We categorise these as: **A.** Short proximal localisation. **B.** pDC-like localisation. **C.** Proximal long localisation.

**Figure S3.**
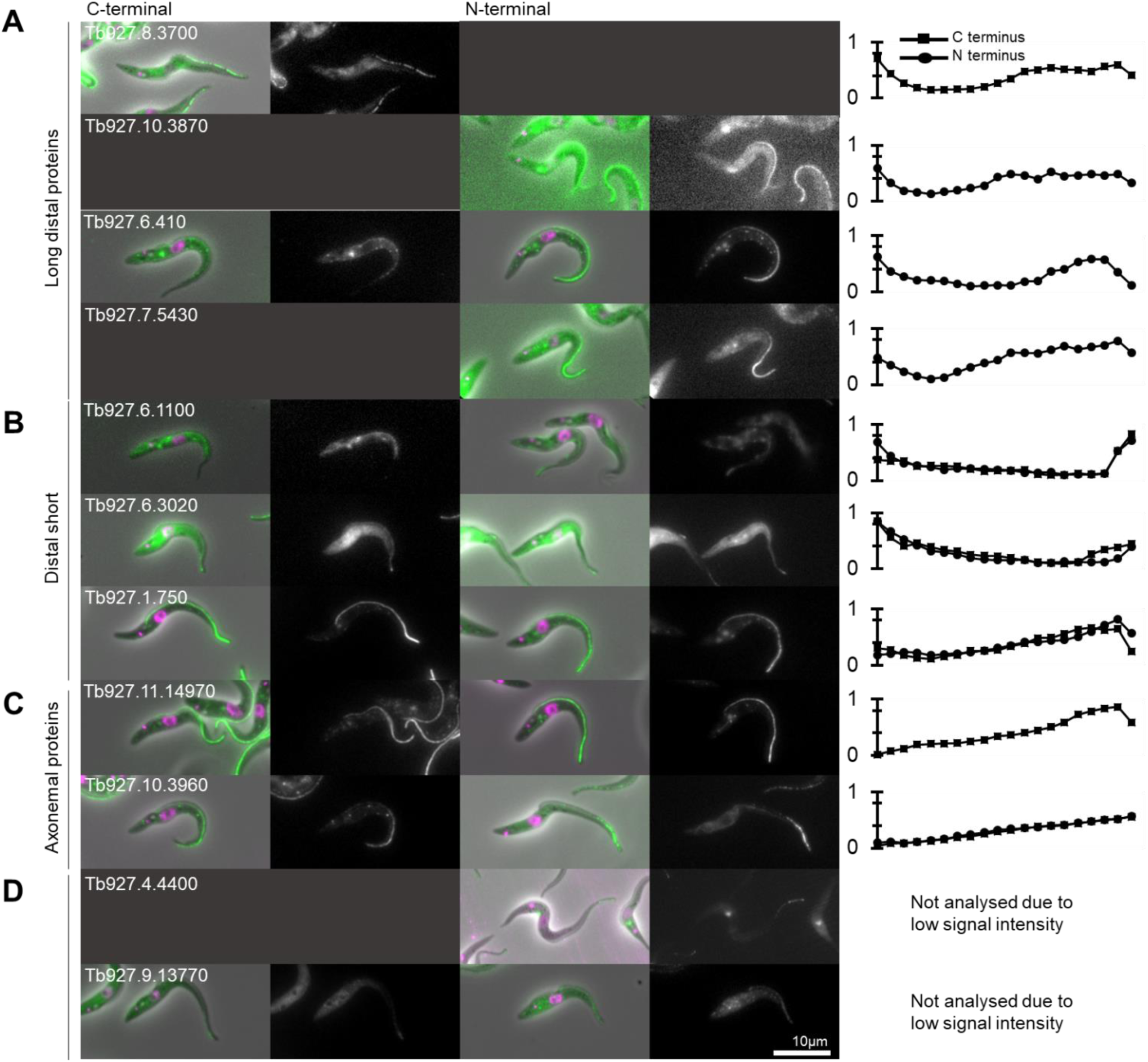
Distal axoneme-specific proteins in *T. brucei* which lack a detectable *L. mexicana* ortholog. In A to D, the first column shows widefield epifluorescence micrographs of mNG fluorescence at the C terminus in three different groups of proximal proteins in *T. brucei*, then the second column shows widefield epifluorescence micrographs of mNG fluorescence at the N terminus in same different groups of proximal proteins in *T. brucei*. Phase contrast (grey), DNA (Hoechst 33342, magenta) and mNG (green) overlay and mNG fluorescence are shown. The third column represent the measure proteins signal intensity along flagellum. Localisations are the same as with N terminal tagging. **A.** Long distal proteins in *T. brucei*. **B.** Distal short proteins in *T. brucei*. **C.** Axonemal proteins in *T. brucei*. **D.** Low signal to determine a specific localisation of proteins in *T. brucei*.

**Figure S4.**
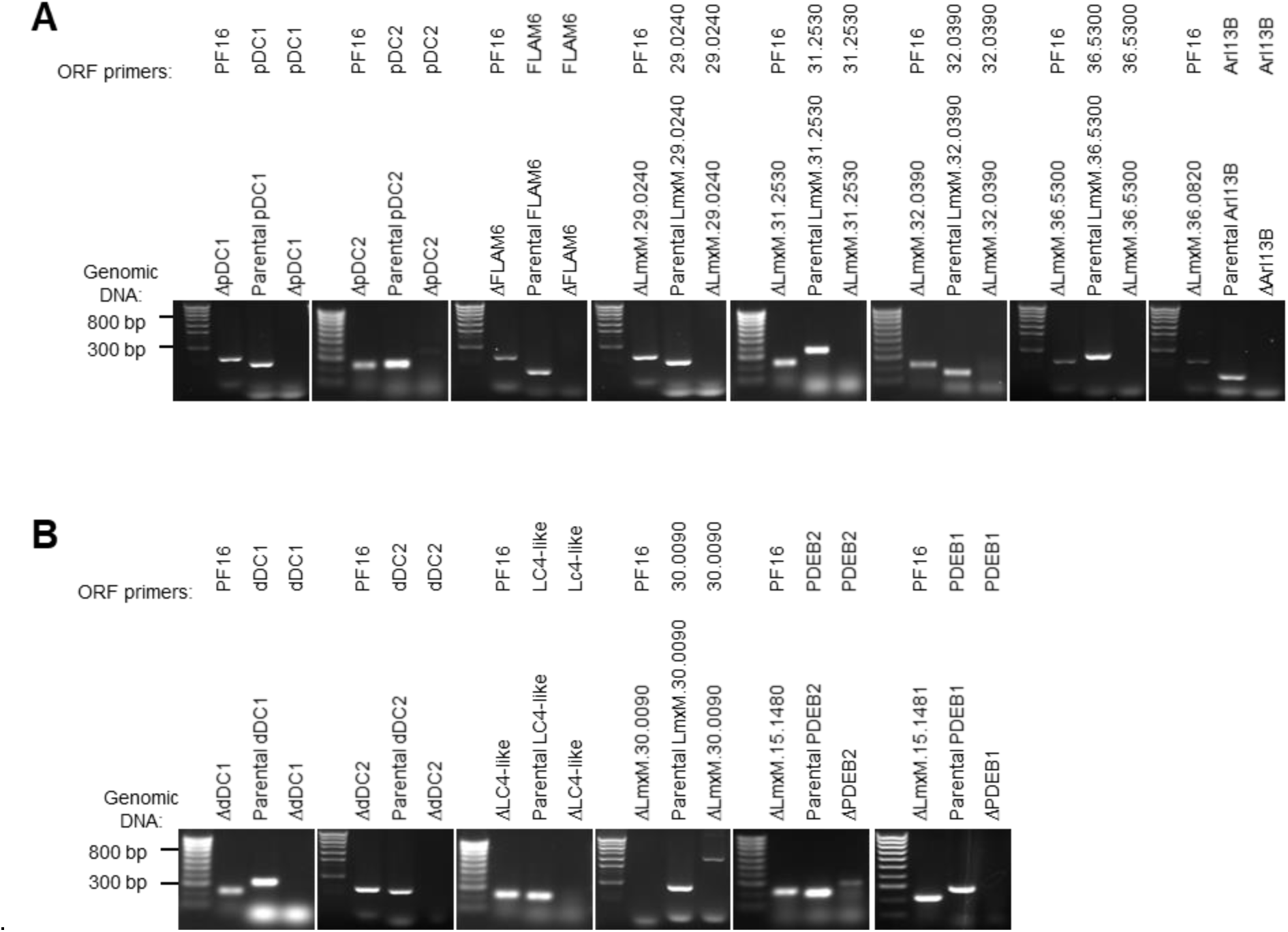
Validation of proximal and distal mutants. Diagnostic PCRs to confirm deletion of both alleles of **A.** Proximal and **B.** Distal proteins in the clonal *L. mexicana* deletion cell lines. For each, gel electrophoresis of PCR products from genomic DNA (gDNA) are shown. Control PCR product from an unaffected open reading frame (ORF), PF16, are shown to confirm presence of deletion mutant gDNA and PCR products from parental gDNA are shown to confirm that the test primers can amplify the dedicated ORF.

**Figure S5.**
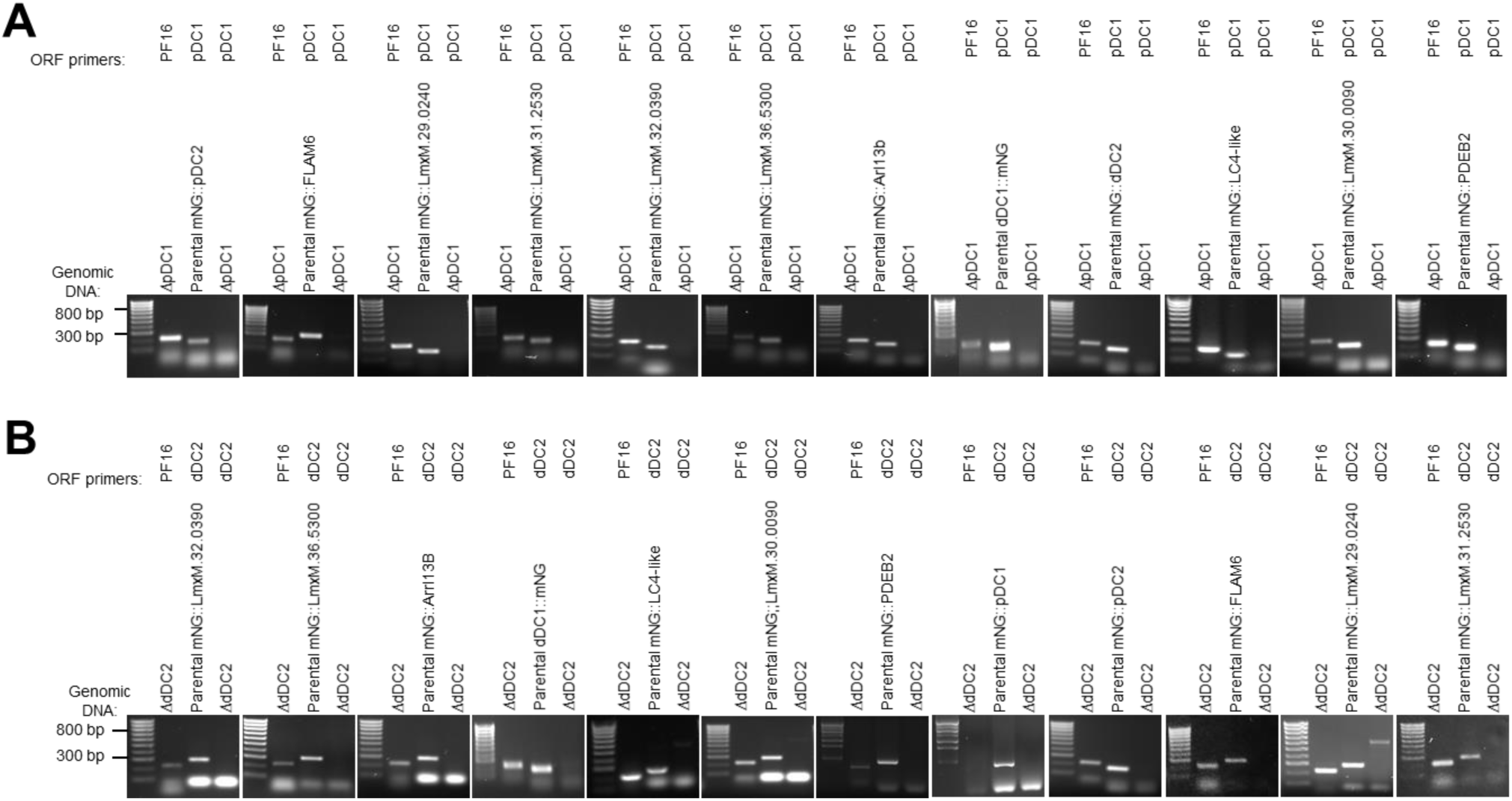
Validation of pDC1 and dDC2 mutants in different tagged cell lines. Diagnostic PCRs to confirm deletion of both alleles of **A.** pDC1 and **B.** dDC2 proteins in the clonal *L. mexicana* deletion cell lines. For each, gel electrophoresis of PCR products from genomic DNA (gDNA) are shown. Control PCR product from an unaffected open reading frame (ORF), PF16, are shown to confirm presence of deletion mutant gDNA and PCR products from parental gDNA are shown to confirm that the test primers can amplify the dedicated ORF.

**Figure S6.**
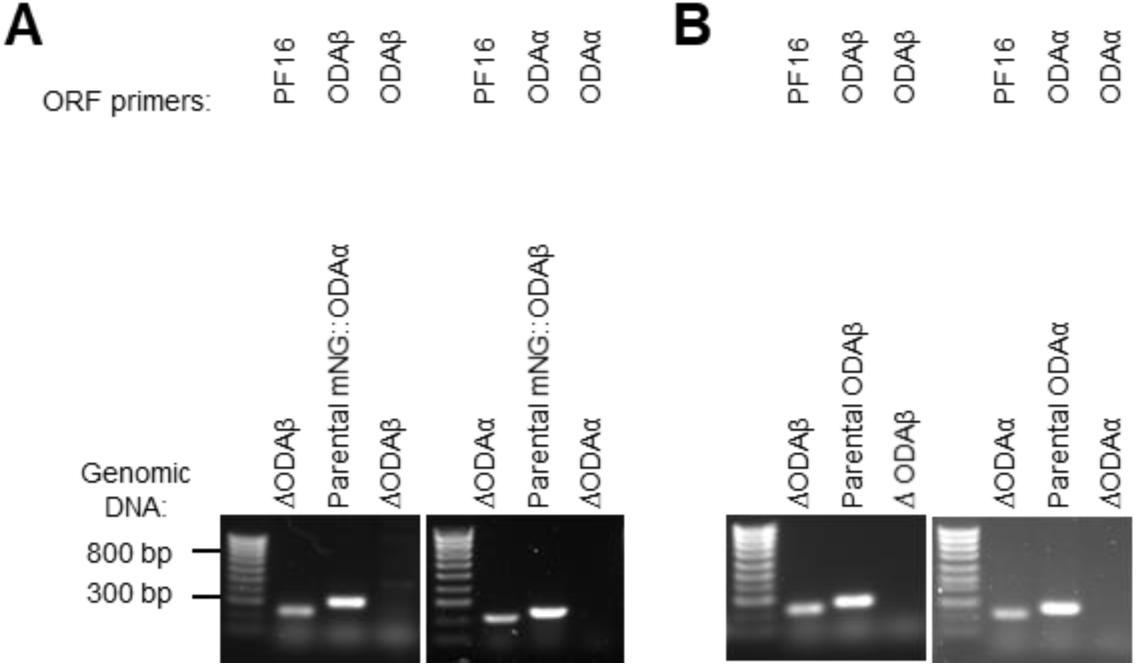
Validation of ODA mutants. Diagnostic PCRs to confirm deletion of both alleles of **A.** ODAβ in tagged mNG::ODAα and ODAα in tagged mNG::ODAβ and **B.** ODAα and ODAβ proteins in the clonal *L. mexicana* deletion cell lines. For each, gel electrophoresis of PCR products from genomic DNA (gDNA) are shown. Control PCR product from an unaffected open reading frame (ORF), PF16, are shown to confirm presence of deletion mutant gDNA and PCR products from parental gDNA are shown to confirm that the test primers can amplify the dedicated ORF.

**Figure S7.**
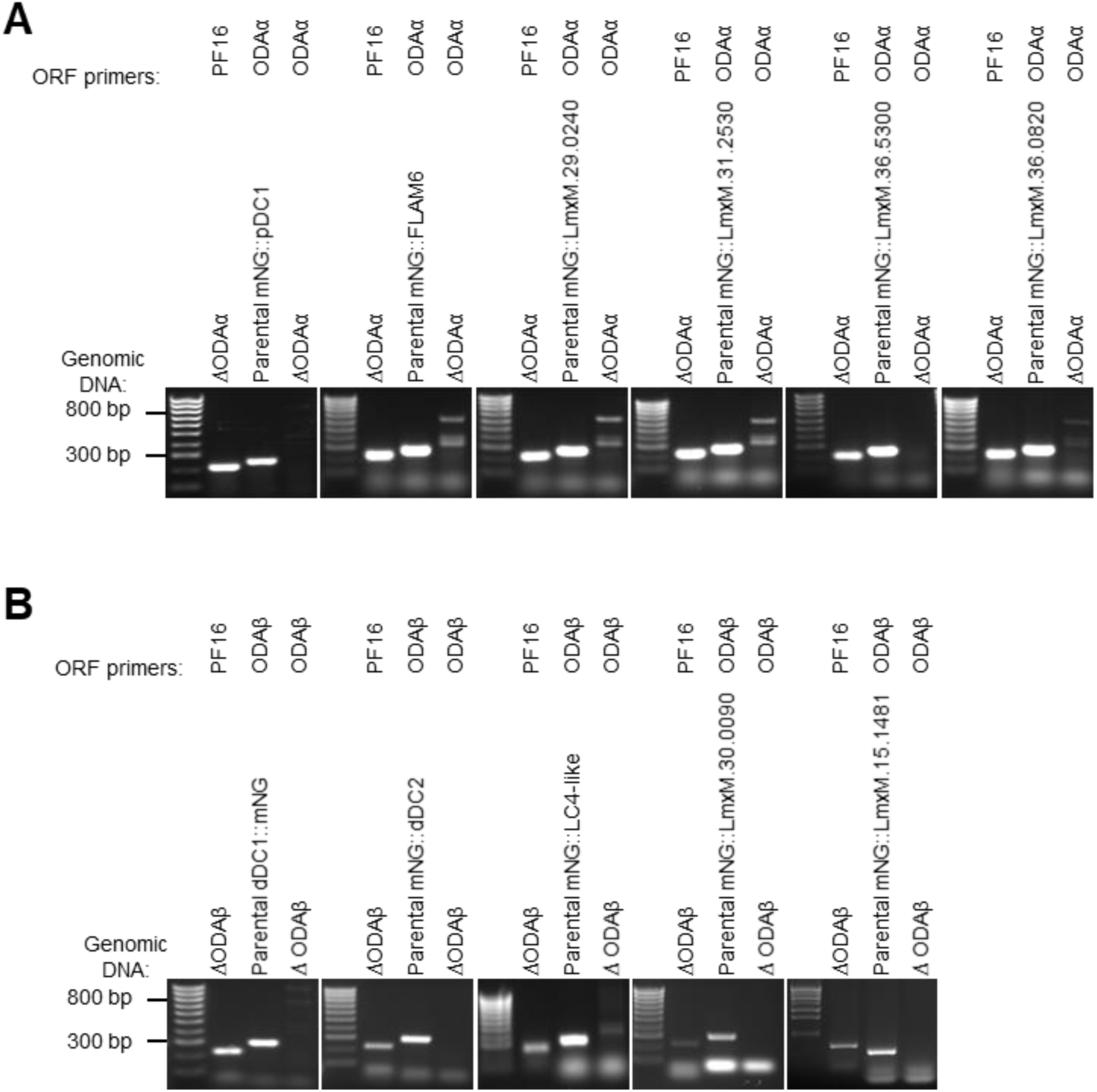
Validation of ODA mutants. Diagnostic PCRs to confirm deletion of both alleles of **A.** ODAα in tagged pDC-like proteins and **B.** ODAβ in tagged dDC-like proteins in the clonal *L. mexicana* deletion cell lines. For each, gel electrophoresis of PCR products from genomic DNA (gDNA) are shown. Control PCR product from an unaffected open reading frame (ORF), PF16, are shown to confirm presence of deletion mutant gDNA and PCR products from parental gDNA are shown to confirm that the test primers can amplify the dedicated ORF.

**Figure S8.**
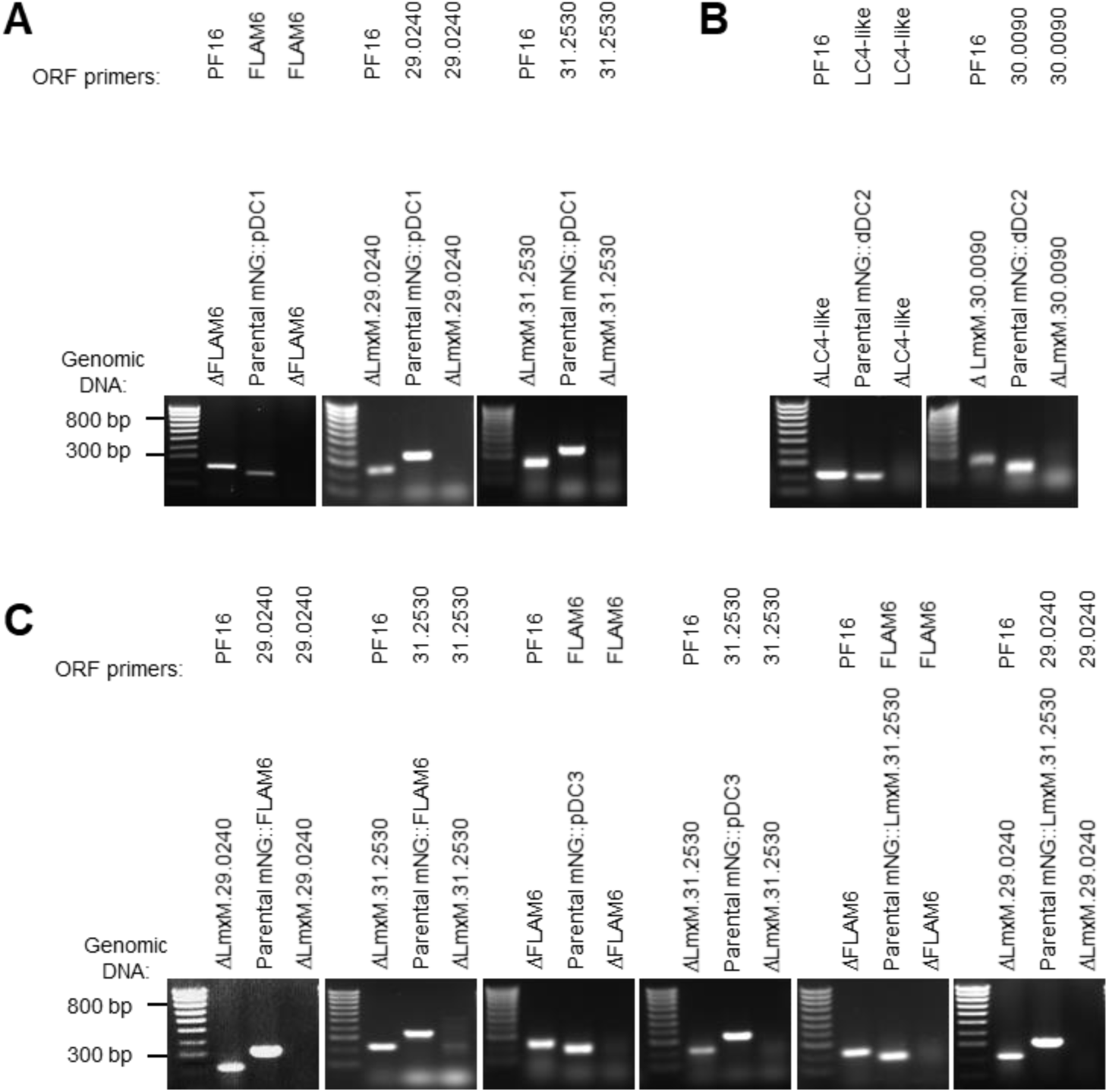
Validation of mutants. Diagnostic PCRs to confirm deletion of both alleles of **A.** pDC-like proteins in tagged pDC1, **B.** dDC-like proteins in tagged dDC2 and **C.** pDC-like proteins in tagged FLAM6, pDC3 and LmxM.31.2530 in the clonal *L. mexicana* deletion cell lines. For each, gel electrophoresis of PCR products from genomic DNA (gDNA) are shown. Control PCR product from an unaffected open reading frame (ORF), PF16, are shown to confirm presence of deletion mutant gDNA and PCR products from parental gDNA are shown to confirm that the test primers can amplify the dedicated ORF.

**Figure S9.**
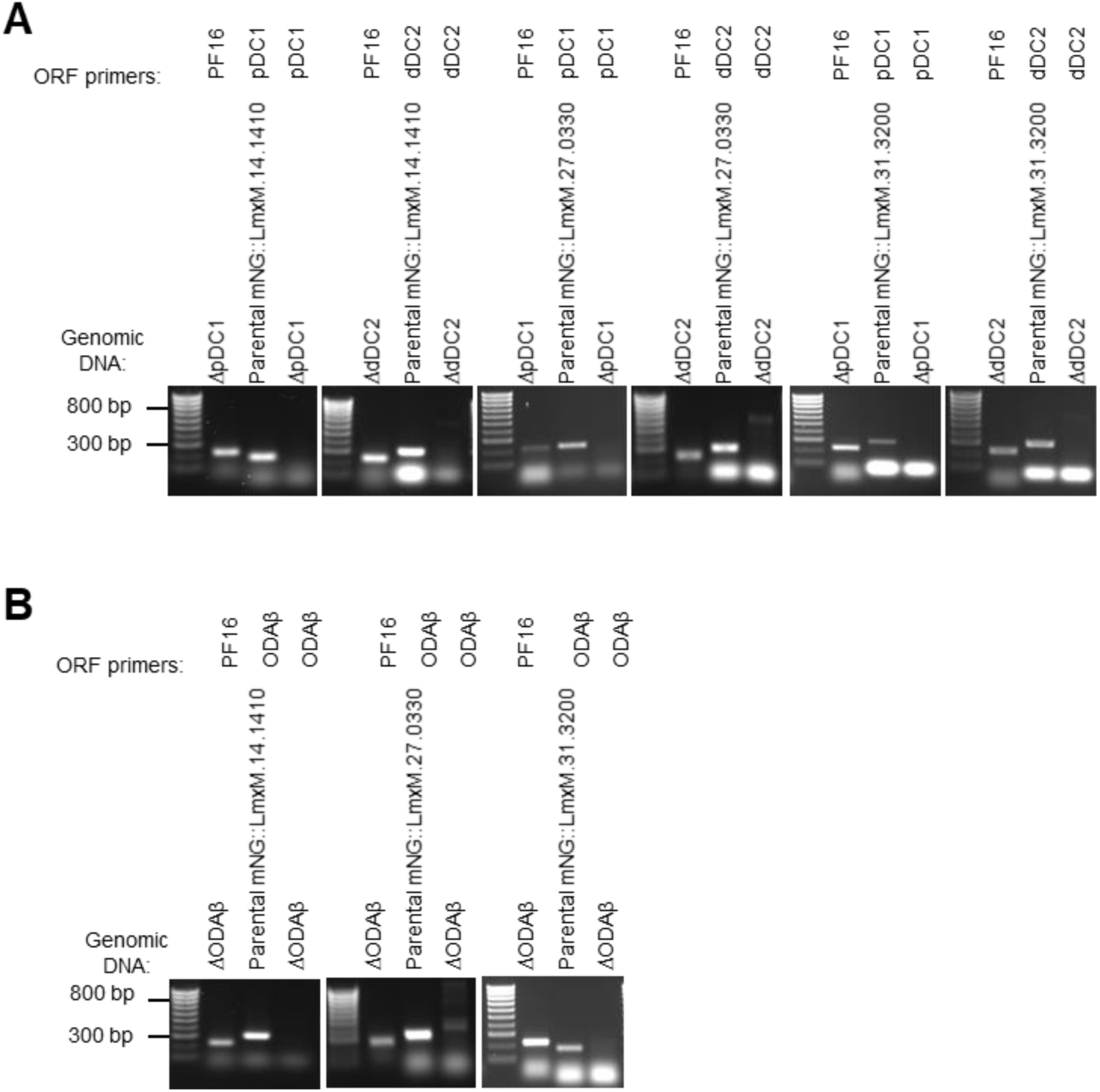
Validation of pDC1, dDC2 and ODAβ mutants in tagged cell lines. Diagnostic PCRs to confirm deletion of both alleles of **A.** pDC1 and dDC2 in tagged of flagellum long tip proteins and **B.** ODAβ in tagged LmxM.14.1410, LmxM.27.0330 and LmxM.31.3200 in the clonal *L. mexicana* deletion cell lines. For each, gel electrophoresis of PCR products from genomic DNA (gDNA) are shown. Control PCR product from an unaffected open reading frame (ORF), PF16, are shown to confirm presence of deletion mutant gDNA and PCR products from parental gDNA are shown to confirm that the test primers can amplify the dedicated ORF.

**Figure S10.**
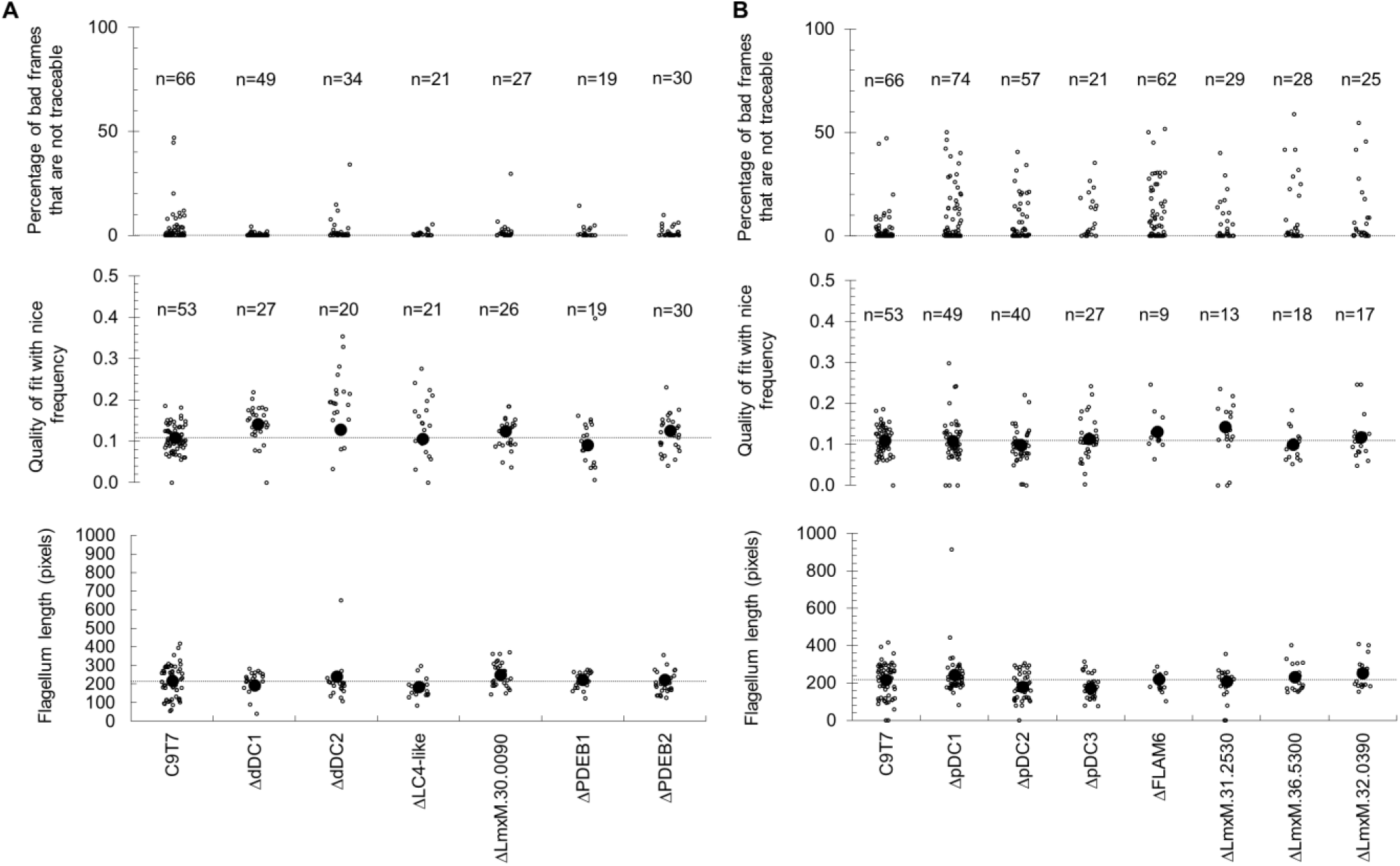
Additional control measures for beat waveform properties for distal and proximal proteins. Graphics represented the percentage of bad frames that are not traceable, the quality of fit with nice frequency and the flagellum length in pixels in **A.** dDC-like and distal enriched PDEs and in **B.** pDC-like and short proteins. n indicates the number of analysed cells.

**Figure S11.**
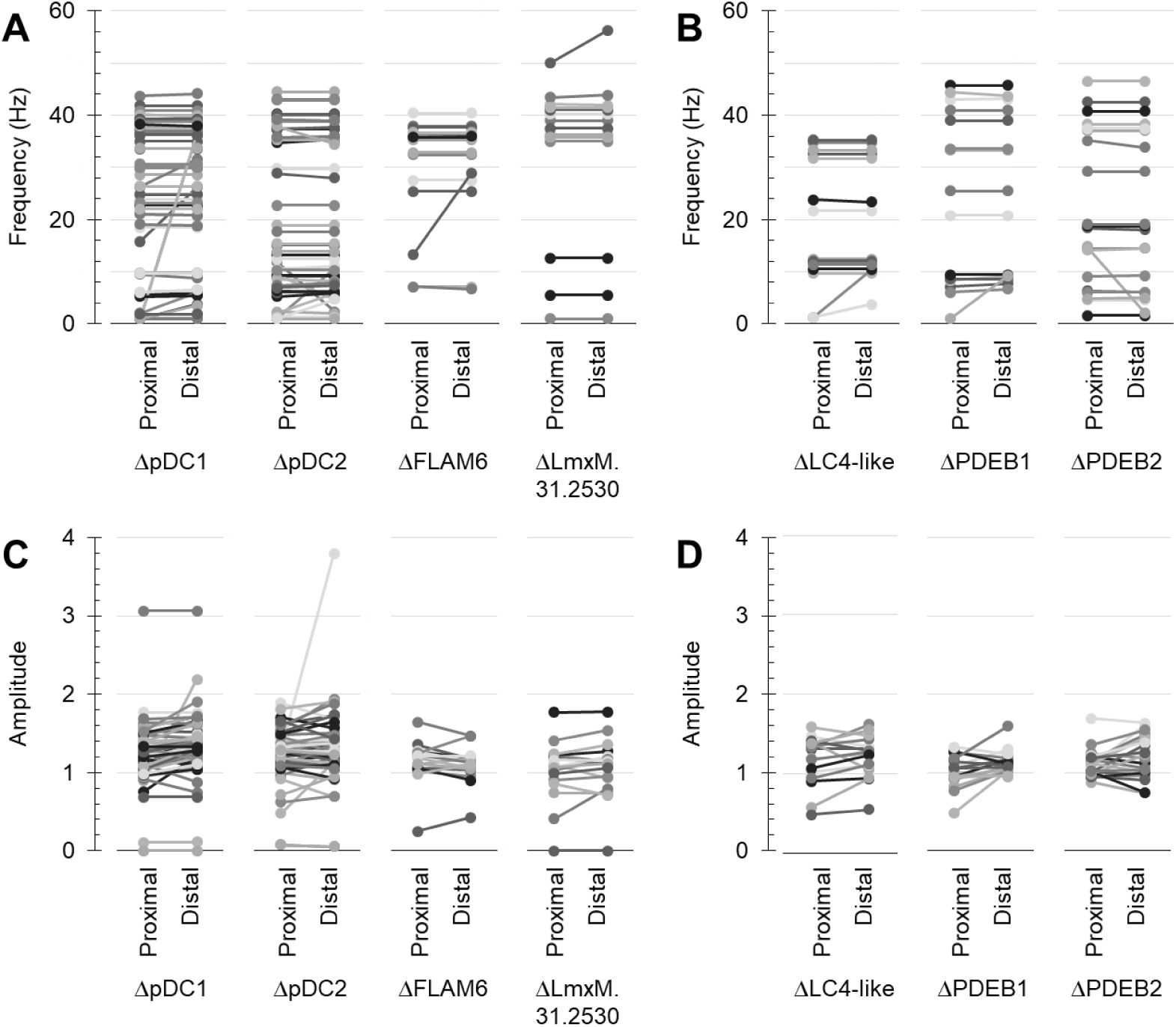
Comparison of dominant beat frequency and amplitude measured from the proximal or distal region of individual flagella, for deletion mutants of proteins with pDC and dDC or distal enriched PDE-like localisations. Every line in each graph represent a single flagellum, with the data points corresponding to beat frequency in the proximal and distal flagella A-C. For deletion mutants of proteins with pDC-like localisation: ΔpDC1, ΔpDC2, ΔFLAM6 and ΔLmxM.31.2530. B-D. For deletion mutants of proteins with distal localisations: ΔLC4-like and ΔPDEB2 or ΔPDEB1. No difference between the proximal and distal flagellum were statistically significant (p > 0.05, two-tailed T test)

**Table S1.**
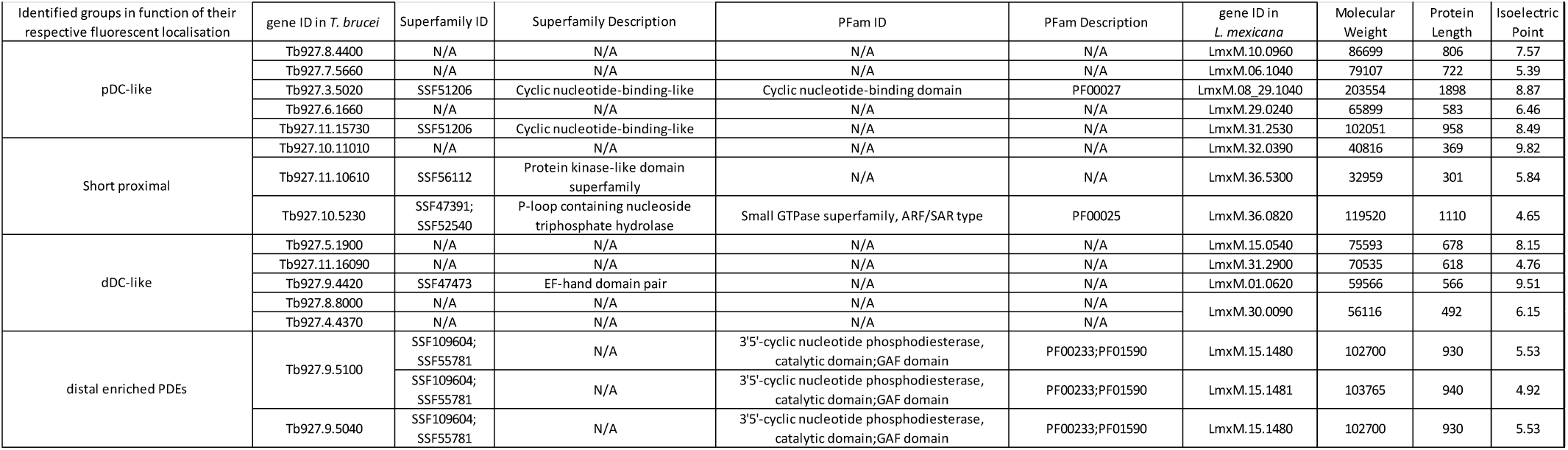
All flagellum proximal and distal proteins in *T. brucei* whose *L. mexicana* ortholog has comparable asymmetric distribution.

**Table S2.**
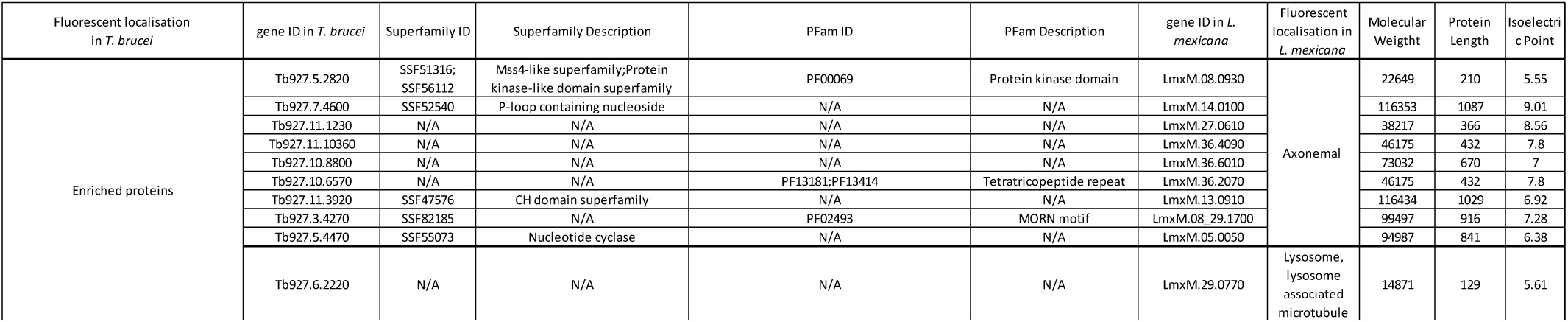
All flagellum proximal and distal axoneme-specific proteins in *T. brucei* whose *L. mexicana* ortholog was not asymmetrically distributed within the flagellum.

**Table S3.** All flagellum proximal and distal proteins in *T. brucei* which lack an *L. mexicana* ortholog.

| Fluorescent localisation in *T. brucei* | gene ID in *T. brucei* | Superfamily ID | Superfamily Description | PFam ID | PFam Description | Molecular Weight | Protein Length | Isoelectric Point |
| --- | --- | --- | --- | --- | --- | --- | --- | --- |
| short proximal | Tb927.1.4340 | N/A | N/A | N/A | N/A | 77650 | 716 | 9.57 |
| short proximal | Tb927.10.7770 | N/A | N/A | N/A | N/A | 25760 | 230 | 10.63 |
| short proximal | Tb927.10.4720 | SSF48403 | Ankyrin repeat-containing domain superfamily | PF13857 | N/A | 39545 | 360 | 4.85 |
| short proximal | Tb927.11.15610 | SSF48452 | Tetratricopeptide-like helical domain superfamily | N/A | N/A | 109790 | 997 | 9.24 |
| short proximal | Tb927.11.5770 | SSF49265 | Fibronectin type III superfamily | N/A | N/A | 222130 | 2030 | 6.1 |
| short proximal | Tb927.11.9650 | N/A | N/A | N/A | N/A | 53172 | 484 | 7.32 |
| pDC-like | Tb927.3.5260 | N/A | N/A | N/A | N/A | 50908 | 469 | 10.54 |
| pDC-like | Tb927.5.1950 | N/A | N/A | N/A | N/A | 66973 | 611 | 8.82 |
| pDC-like | Tb927.6.3410 | N/A | N/A | N/A | N/A | 34704 | 311 | 8.38 |
| pDC-like | Tb927.11.15740 | N/A | N/A | N/A | N/A | 60161 | 542 | 11.36 |
| pDC-like | Tb927.3.1200 | SSF47473 | EF-hand domain pair | N/A | N/A | 28693 | 263 | 9.85 |
| long proximal | Tb927.11.5790 | N/A | N/A | N/A | N/A | 30645 | 273 | 10 |
| long proximal | Tb927.8.1280 | N/A | N/A | N/A | N/A | 153905 | 1393 | 9.01 |
| long proximal | Tb927.9.2075 | N/A | N/A | N/A | N/A | 302504 | 2843 | 4.2 |
| long proximal | Tb927.9.7100 | N/A | N/A | N/A | N/A | 25626 | 246 | 6.13 |
| long proximal | Tb927.9.9300 | N/A | N/A | N/A | N/A | 54512 | 482 | 9.69 |
| long proximal | Tb927.6.2120 | N/A | N/A | N/A | N/A | 27386 | 250 | 10.45 |
| long proximal | Tb927.8.5300 | N/A | N/A | N/A | N/A | 63847 | 562 | 10.59 |
| long distal | Tb927.10.3870 | SSF101908;<br>SSF47473;<br>SSF50978 | EF-hand domain pair;WD40-repeat-containing domain superfamily | PF00400 | WD40 repeat | 237134 | 2151 | 7.79 |
| long distal | Tb927.7.5430 | N/A | N/A | N/A | N/A | 98743 | 906 | 9.22 |
| long distal | Tb927.8.3700 | N/A | N/A | N/A | N/A | 161957 | 1486 | 8.65 |
| long distal | Tb927.6.410 | SSF52058 | N/A | N/A | N/A | 93966 | 858 | 4.93 |
| long distal | Tb927.7.5430 | N/A | N/A | N/A | N/A | 98743 | 906 | 9.22 |
| short distal | Tb927.6.1100 | N/A | N/A | N/A | N/A | 53686 | 492 | 7.6 |
| short distal | Tb927.6.3020 | N/A | N/A | N/A | N/A | 32217 | 288 | 10.23 |
| short distal | Tb927.1.750 | N/A | N/A | N/A | N/A | 132691 | 1215 | 6.86 |
| axonemal | Tb927.10.3960 | N/A | N/A | N/A | N/A | 17598 | 155 | 4.69 |
| axonemal | Tb927.11.14970 | N/A | N/A | N/A | N/A | 93163 | 823 | 5.13 |
|  | Tb927.4.4400 | SSF56104 | N/A | PF03770 | Inositol polyphosphate kinase | 82630 | 756 | 5.63 |
|  | Tb927.9.13770 | N/A | N/A | PF04683 | Proteasomal ubiquitin receptor Rpn13/ADRM1 | 31304 | 279 | 5.38 |

**Table S4.**
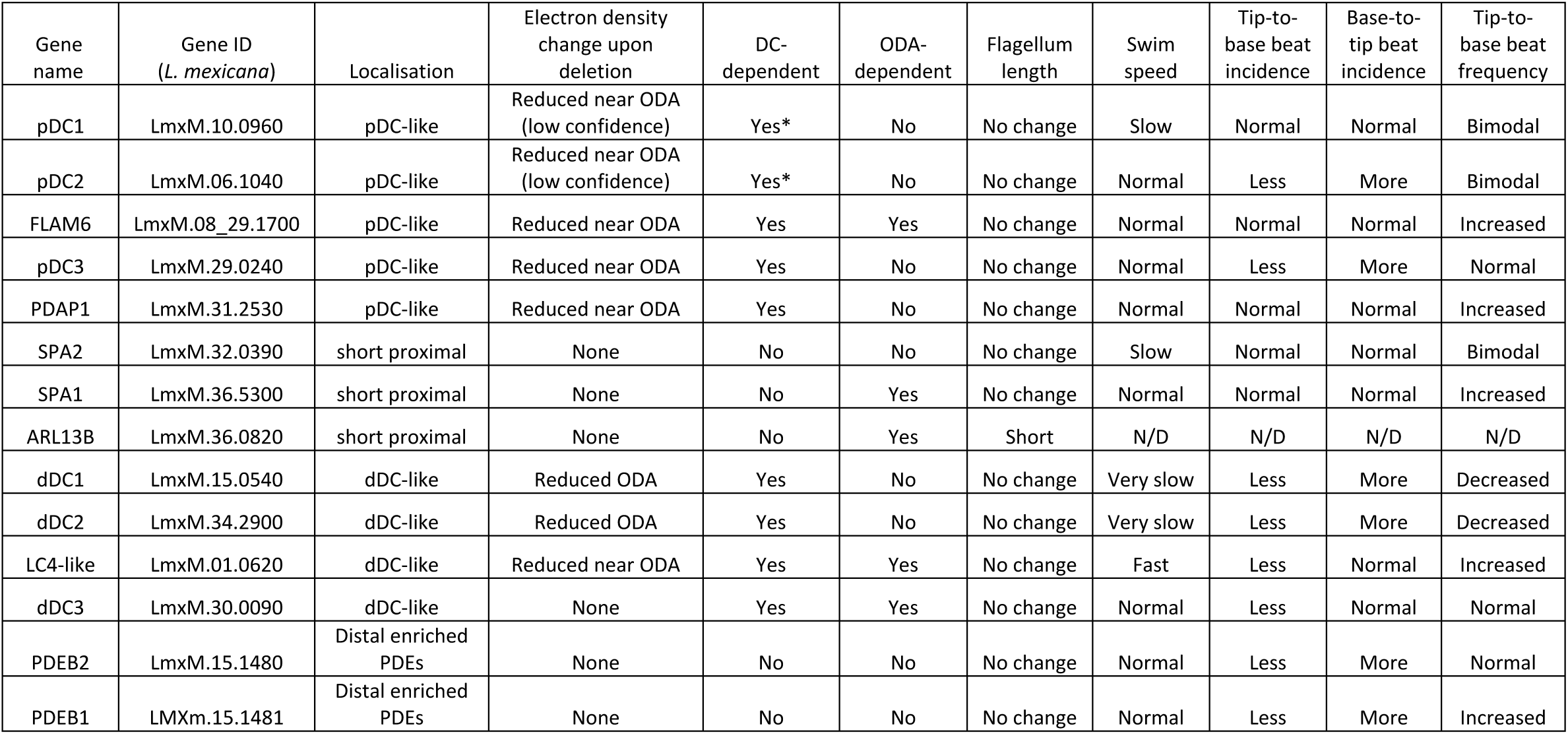
Qualitative summary of protein localisation and deletion mutant phenotypes.

## References

1. Edwards, B. F. L. et al. Direction of flagellum beat propagation is controlled by proximal/distal outer dynein arm asymmetry. Proc Natl Acad Sci U S A 115, E7341–E7350 (2018).

2. Gadelha, C., Wickstead, B. & Gull, K. Flagellar and ciliary beating in trypanosome motility. Cell Motility 64, 629–643 (2007).

3. Shiba, K., Shibata, D. & Inaba, K. Autonomous changes in the swimming direction of sperm in the gastropod *Strombus luhuanus*. Journal of Experimental Biology jeb.095398 (2013) doi:10.1242/jeb.095398.

4. Yang, Y. & Lu, X. Drosophila Sperm Motility in the Reproductive Tract. Biol Reprod 84, 1005– 1015 (2011).

5. Baron, D. M., Kabututu, Z. P. & Hill, K. L. Stuck in reverse: loss of LC1 in *Trypanosoma brucei* disrupts outer dynein arms and leads to reverse flagellar beat and backward movement. Journal of Cell Science 120, 1513–1520 (2007).

6. Wheeler, R. J. Use of chiral cell shape to ensure highly directional swimming in trypanosomes. PLOS Computational Biology 13, e1005353 (2017).

7. Santi-Rocca, J. et al. Imaging intraflagellar transport in trypanosomes. Methods Cell Biol 127, 487–508 (2015).

8. Beneke, T. et al. A CRISPR Cas9 high-throughput genome editing toolkit for kinetoplastids. R Soc Open Sci 4, 170095 (2017).

9. Beneke, T. et al. Genetic dissection of a Leishmania flagellar proteome demonstrates requirement for directional motility in sand fly infections. PLoS Pathog 15, e1007828 (2019).

10. Rotureau, B., Ooi, C.-P., Huet, D., Perrot, S. & Bastin, P. Forward motility is essential for trypanosome infection in the tsetse fly. Cell Microbiol 16, 425–433 (2014).

11. Brokaw, C. J. & Kamiya, R. Bending patterns of *Chlamydomonas* flagella: IV. Mutants with defects in inner and outer dynein arms indicate differences in dynein arm function. Cell Motil. Cytoskeleton 8, 68–75 (1987).

12. Bui, K. H., Sakakibara, H., Movassagh, T., Oiwa, K. & Ishikawa, T. Molecular architecture of inner dynein arms in situ in Chlamydomonas reinhardtii flagella. J Cell Biol 183, 923–932 (2008).

13. Takada, S., Wilkerson, C. G., Wakabayashi, K., Kamiya, R. & Witman, G. B. The Outer Dynein Arm-Docking Complex: Composition and Characterization of a Subunit (Oda1) Necessary for Outer Arm Assembly. Mol Biol Cell 13, 1015–1029 (2002).

14. Caspary, T., Larkins, C. E. & Anderson, K. V. The Graded Response to Sonic Hedgehog Depends on Cilia Architecture. Developmental Cell 12, 767–778 (2007).

15. Cevik, S. et al. Joubert syndrome Arl13b functions at ciliary membranes and stabilizes protein transport in Caenorhabditis elegans. J Cell Biol 188, 953–969 (2010).

16. Duldulao, N. A., Lee, S. & Sun, Z. Cilia localization is essential for in vivo functions of the Joubert syndrome protein Arl13b/Scorpion. Development 136, 4033–4042 (2009).

17. Li, Y., Wei, Q., Zhang, Y., Ling, K. & Hu, J. The small GTPases ARL-13 and ARL-3 coordinate intraflagellar transport and ciliogenesis. J Cell Biol 189, 1039–1051 (2010).

18. Lu, H. et al. A function for the Joubert syndrome protein Arl13b in ciliary membrane extension and ciliary length regulation. Dev Biol 397, 225–236 (2015).

19. Ishikawa, H. & Marshall, W. F. Ciliogenesis: building the cell’s antenna. Nat Rev Mol Cell Biol 12, 222–234 (2011).

20. Cantagrel, V. et al. Mutations in the cilia gene ARL13B lead to the classical form of Joubert syndrome. Am J Hum Genet 83, 170–179 (2008).

21. Zhang, Y. et al. The unusual flagellar-targeting mechanism and functions of the trypanosome ortholog of the ciliary GTPase Arl13b. J Cell Sci 131, jcs219071 (2018).

22. Cevik, S. et al. Active Transport and Diffusion Barriers Restrict Joubert Syndrome-Associated ARL13B/ARL-13 to an Inv-like Ciliary Membrane Subdomain. PLoS Genet 9, e1003977 (2013).

23. Oberholzer, M. et al. The Trypanosoma brucei cAMP phosphodiesterases TbrPDEBl and TbrPDEB2: flagellar enzymes that are essential for parasite virulence. The FASEB Journal 21, 720–731 (2007).

24. Oberholzer, M., Saada, E. A. & Hill, K. L. Cyclic AMP Regulates Social Behavior in African Trypanosomes. mBio 6, e01954–14 (2015).

25. Nlend, M.-C. et al. Calcium-mediated, purinergic stimulation and polarized localization of calcium-sensitive adenylyl cyclase isoforms in human airway epithelia. FEBS Lett 581, 3241–3246 (2007).

26. Balbach, M., Beckert, V., Hansen, J. N. & Wachten, D. Shedding light on the role of cAMP in mammalian sperm physiology. Mol Cell Endocrinol 468, 111–120 (2018).

27. Aoki, F., Sakai, S. & Kohmoto, K. Regulation of flagellar bending by cAMP and Ca2+ in hamster sperm. Mol Reprod Dev 53, 77–83 (1999).

28. Beltrán, C., Zapata, O. & Darszon, A. Membrane Potential Regulates Sea Urchin Sperm Adenylylcyclase. Biochemistry 35, 7591–7598 (1996).

29. Hyams, J. S. & Borisy, G. G. Isolated flagellar apparatus of Chlamydomonas: characterization of forward swimming and alteration of waveform and reversal of motion by calcium ions in vitro. Journal of Cell Science 33, 235–253 (1978).

30. Differential regulation of Paramecium ciliary motility by cAMP and cGMP. J Cell Biol 106, 1615–1623 (1988).

31. Mukhopadhyay, A. G. & Dey, C. S. Role of calmodulin and calcineurin in regulating flagellar motility and wave polarity in Leishmania. Parasitol Res 116, 3221–3228 (2017).

32. Pivato, M. & Ballottari, M. Chlamydomonas reinhardtii cellular compartments and their contribution to intracellular calcium signalling. J Exp Bot 72, 5312–5335 (2021).

33. Harz, H. & Hegemann, P. Rhodopsin-regulated calcium currents in Chlamydomonas. Nature 351, 489–491 (1991).

34. Pazour, G. J., Sineshchekov, O. A. & Witman, G. B. Mutational analysis of the phototransduction pathway of Chlamydomonas reinhardtii. J Cell Biol 131, 427–440 (1995).

35. Mukhopadhyay, A. G. & Dey, C. S. Effect of inhibition of axonemal dynein ATPases on the regulation of flagellar and ciliary waveforms in Leishmania parasites. Mol Biochem Parasitol 225, 27–37 (2018).

36. Subota, I. et al. Proteomic Analysis of Intact Flagella of Procyclic Trypanosoma brucei Cells Identifies Novel Flagellar Proteins with Unique Sub-localization and Dynamics. Mol Cell Proteomics 13, 1769–1786 (2014).

37. Shaw, S. et al. Flagellar cAMP signaling controls trypanosome progression through host tissues. Nat Commun 10, 803 (2019).

38. Billington, K. et al. Genome-wide subcellular protein map for the flagellate parasite Trypanosoma brucei. Nat Microbiol 8, 533–547 (2023).

39. Jackson, A. P. Evolutionary consequences of a large duplication event in Trypanosoma brucei: Chromosomes 4 and 8 are partial duplicons. BMC Genomics 8, 432 (2007).

40. Song, P., Dudinsky, L., Fogerty, J., Gaivin, R. & Perkins, B. D. Arl13b Interacts With Vangl2 to Regulate Cilia and Photoreceptor Outer Segment Length in Zebrafish. Invest Ophthalmol Vis Sci 57, 4517–4526 (2016).

41. Parisi, M. A. Clinical and molecular features of Joubert syndrome and related disorders. American Journal of Medical Genetics Part C: Seminars in Medical Genetics 151C, 326–340 (2009).

42. Alkanderi, S., et al. *ARL3* Mutations Cause Joubert Syndrome by Disrupting Ciliary Protein Composition. The American Journal of Human Genetics 103, 612–620 (2018).

43. Gadelha, C., Wickstead, B., McKean, P. G. & Gull, K. Basal body and flagellum mutants reveal a rotational constraint of the central pair microtubules in the axonemes of trypanosomes. J Cell Sci 119, 2405–2413 (2006).

44. Gluenz, E., Wheeler, R. J., Hughes, L. & Vaughan, S. Scanning and three-dimensional electron microscopy methods for the study of Trypanosoma brucei and Leishmania mexicana flagella. Methods Cell Biol 127, 509–542 (2015).

45. Chiurillo, M. A., Ahmed, M., González, C., Raja, A. & Lander, N. Gene editing of putative cAMP and Ca2+-regulated proteins using an efficient cloning-free CRISPR/Cas9 system in Trypanosoma cruzi. Journal of Eukaryotic Microbiology 70, e12999 (2023).

46. Fowkes, M. E. & Mitchell, D. R. The Role of Preassembled Cytoplasmic Complexes in Assembly of Flagellar Dynein Subunits. Mol Biol Cell 9, 2337–2347 (1998).

47. Mitchell, D. R. & Rosenbaum, J. L. A motile Chlamydomonas flagellar mutant that lacks outer dynein arms. Journal of Cell Biology 100, 1228–1234 (1985).

48. Kamiya, R. Mutations at twelve independent loci result in absence of outer dynein arms in Chylamydomonas reinhardtii. Journal of Cell Biology 107, 2253–2258 (1988).

49. Bui, K. H., Yagi, T., Yamamoto, R., Kamiya, R. & Ishikawa, T. Polarity and asymmetry in the arrangement of dynein and related structures in the Chlamydomonas axoneme. J Cell Biol 198, 913–925 (2012).

50. Fliegauf, M. et al. Mislocalization of DNAH5 and DNAH9 in respiratory cells from patients with primary ciliary dyskinesia. Am J Respir Crit Care Med 171, 1343–1349 (2005).

51. Panizzi, J. R. et al. CCDC103 mutations cause primary ciliary dyskinesia by disrupting assembly of ciliary dynein arms. Nat Genet 44, 714–719 (2012).

52. Hoops, H. J. & Witman, G. B. Outer doublet heterogeneity reveals structural polarity related to beat direction in Chlamydomonas flagella. J Cell Biol 97, 902–908 (1983).

53. Dean, A. B. & Mitchell, D. R. Late steps in cytoplasmic maturation of assembly-competent axonemal outer arm dynein in Chlamydomonas require interaction of ODA5 and ODA10 in a complex. Mol Biol Cell 26, 3596–3605 (2015).

54. Hagen, K. D. et al. Novel structural components of the ventral disc and lateral crest in Giardia intestinalis. PLoS Negl Trop Dis 5, e1442 (2011).

55. Jumper, J. et al. Highly accurate protein structure prediction with AlphaFold. Nature 596, 583–589 (2021).

56. Mirdita, M. et al. ColabFold: making protein folding accessible to all. Nat Methods 19, 679–682 (2022).

57. Wheeler, R. J. A resource for improved predictions of Trypanosoma and Leishmania protein three-dimensional structure. PLOS ONE 16, e0259871 (2021).

58. Pugacheva, E. N., Jablonski, S. A., Hartman, T. R., Henske, E. P. & Golemis, E. A. HEF1-dependent Aurora A activation induces disassembly of the primary cilium. Cell 129, 1351–1363 (2007).

59. Ran, J., Yang, Y., Li, D., Liu, M. & Zhou, J. Deacetylation of α-tubulin and cortactin is required for HDAC6 to trigger ciliary disassembly. Sci Rep 5, 12917 (2015).

60. Sherwin, T., Schneider, A., Sasse, R., Seebeck, T. & Gull, K. Distinct localization and cell cycle dependence of COOH terminally tyrosinolated alpha-tubulin in the microtubules of Trypanosoma brucei brucei. Journal of Cell Biology 104, 439–446 (1987).

61. Sasse, R. & Gull, K. Tubulin post-translational modifications and the construction of microtubular organelles in Trypanosoma brucei. J Cell Sci 90 **(Pt** **4**), 577–589 (1988).

62. Sloboda, R. D. Posttranslational Protein Modifications in Cilia and Flagella. in Methods in Cell Biology vol. 94 347–363 (Elsevier, 2009).

63. Bastin, P., Ellis, K., Kohl, L. & Gull, K. Flagellum ontogeny in trypanosomes studied via an inherited and regulated RNA interference system. J Cell Sci 113 **(Pt** **18**), 3321–3328 (2000).

64. Wheeler, R. J., Sunter, J. D. & Gull, K. Flagellar pocket restructuring through the *Leishmania* life cycle involves a discrete flagellum attachment zone. Journal of Cell Science jcs.183152 (2016) doi:10.1242/jcs.183152.

65. Kohl, L., Robinson, D. & Bastin, P. Novel roles for the flagellum in cell morphogenesis and cytokinesis of trypanosomes. EMBO J 22, 5336–5346 (2003).

66. Madden, T. The BLAST Sequence Analysis Tool. in The NCBI Handbook [Internet] (National Center for Biotechnology Information (US), 2003).

67. Billington, K., et al. TrypTag: Genome-wide subcellular protein localisation in Trypanosoma brucei. Zenodo 10.5281/zenodo.7719295 (2022).

68. Dean, S. et al. A toolkit enabling efficient, scalable and reproducible gene tagging in trypanosomatids. Open Biol 5, 140197 (2015).

69. Collins, T. J. ImageJ for microscopy. Biotechniques 43, 25–30 (2007).

70. Automated identification of flagella from videomicroscopy via the medial axis transform | Scientific Reports. https://www.nature.com/articles/s41598-019-41459-9.

71. Reynolds, E. S. THE USE OF LEAD CITRATE AT HIGH pH AS AN ELECTRON-OPAQUE STAIN IN ELECTRON MICROSCOPY. J Cell Biol 17, 208–212 (1963).

72. Gadelha, A. P. R., Cunha-e-Silva, N. L. & de Souza, W. Assembly of the Leishmania amazonensis flagellum during cell differentiation. Journal of Structural Biology 184, 280–292 (2013).

